# TF-Prioritizer: a java pipeline to prioritize condition-specific transcription factors

**DOI:** 10.1101/2022.10.19.512881

**Authors:** Markus Hoffmann, Nico Trummer, Leon Schwartz, Jakub Jankowski, Hye Kyung Lee, Lina-Liv Willruth, Olga Lazareva, Kevin Yuan, Nina Baumgarten, Florian Schmidt, Jan Baumbach, Marcel H. Schulz, David B. Blumenthal, Lothar Hennighausen, Markus List

## Abstract

**Background:** Eukaryotic gene expression is controlled by cis-regulatory elements (CREs), including promoters and enhancers, which are bound by transcription factors (TFs). Differential expression of TFs and their binding affinity at putative CREs determine tissue- and developmental-specific transcriptional activity. Consolidating genomic data sets can offer further insights into the accessibility of CREs, TF activity, and, thus, gene regulation. However, the integration and analysis of multi-modal data sets are hampered by considerable technical challenges. While methods for highlighting differential TF activity from combined chromatin state data (e.g., ChIP-seq, ATAC-seq, or DNase-seq) and RNA-seq data exist, they do not offer convenient usability, have limited support for large-scale data processing, and provide only minimal functionality for visually interpreting results.

**Results:** We developed TF-Prioritizer, an automated pipeline that prioritizes condition-specific TFs from multi-modal data and generates an interactive web report. We demonstrated its potential by identifying known TFs along with their target genes, as well as previously unreported TFs active in lactating mouse mammary glands. Additionally, we studied a variety of ENCODE data sets for cell lines K562 and MCF-7, including twelve histone modification ChIP-seq as well as ATAC-seq and DNase-seq datasets, where we observe and discuss assay-specific differences.

**Conclusion:** TF-Prioritizer accepts ATAC-seq, DNase-seq, or ChIP-seq and RNA-seq data as input and identifies TFs with differential activity, thus offering an understanding of genome-wide gene regulation, potential pathogenesis, and therapeutic targets in biomedical research.

## INTRODUCTION

Understanding how genes are regulated remains a major research focus of molecular biology and genetics [1]. In eukaryotes, gene expression is controlled by cis-regulatory elements (CREs) such as promoters, enhancers, or suppressors, which are bound by transcription factors (TFs) promoting or repressing transcriptional activity depending on their accessibility [2]. TFs play an important role not only in development and physiology but also in diseases, e.g., it is known that at least a third of all known human developmental disorders are associated with deregulated TF activity and mutations [3–5]. An in-depth investigation of TF regulation could help to gain deeper insights into the gene-regulatory balance found in normal physiology. Since most complex diseases involve aberrant gene regulation, a detailed understanding of this mechanism is a prerequisite to developing targeted therapies [6,7]. This is a daunting task, as multiple genes in eukaryotic genomes may affect the disease, each of which is possibly controlled by candidate CREs.

TF ChIP-seq experiments are the gold standard for identifying and understanding condition-specific TF-binding at a nucleotide level. However, since there are approximately 1,500 active TFs in humans [8] and about 1,000 in mice [9], and additionally considering the need to establish TF patterns separately for each tissue and physiological condition, this approach is logistically prohibitive. Alternatively, histone modification (HM) ChIP-seq, but also ATAC-seq and DNAse-seq offer a broader view of the chromatin state due to their individual capability (i.e., ChIP-seq through identifying protein-DNA interactions, ATAC-seq detects open chromatin regions via Tn5 transposase cuttings, and DNAse-seq maps accessible chromatin sites by digesting chromatin with DNase I) to highlight open chromatin regions aligned with active genes, hence allowing the identification of condition-specific CREs [10]. Computational methods can then be used to prioritize TFs likely binding to these CREs, leading to hypotheses and defining the most promising TF ChIP-seq experiments. This narrows the scope of TF ChIP-seq experiments needed to confirm working hypotheses about gene regulation [11–13].

Several general approaches have been proposed to identify key TFs that are responsible for gene regulation. Among them, e.g., (1) a basic coexpression or mutual information analysis of TFs and their target genes combined with computational binding site predictions [14]. (2) Some tools use a combination of TF ChIP-seq data - providing genome-wide information about the exact locations of TF binding - with predicted target genes that can enhance co-expression analyses [15]. (3) Other tools employ a combination of genome-wide chromatin accessibility (e.g., HM ChIP-seq data) or activity information, putative TF binding sites, and gene expression data. This combination can be powerful in determining key TF players and is used by the state-of-the-art tool diffTF [16]. Most of the proposed approaches require substantial preprocessing, computational knowledge, adjustment of the method to a new use case (e.g., more than two conditions and/or time-series data), and manual evaluation of the results (e.g., manual search and visualization for TF ChIP-seq data to provide experimental evidence for the predictions). Hence, to streamline this process, we present TF-Prioritizer, a java pipeline to prioritize TFs that show condition-specific changes in their activity. TF-Prioritizer falls into the third category of the previously described approaches and automates several time-consuming steps, including data processing, TF affinity analysis, machine learning predicting relationships of CREs to target genes, prioritization of relevant TFs, data visualization, and visual experimental validation of the findings using public TF ChIP-seq data (i.e., ChIP-Atlas [17]).

Figure 1 depicts a general overview of the pipeline. TF-Prioritizer accepts two types of input data: i) histone modification peak ChIP-seq/ATAC-seq/DNase-seq data indicating accessible regulatory regions showing differential activity (peak data is typically generated by MACS2 [18]), and ii) gene expression data from RNA-seq, which allows the identification of differentially expressed genes that are potentially regulated by TFs at specific time points or physiological condition. If peaks from ATAC-seq or DNase-seq were provided, we generate footprints (i.e., specific regions of the peaks within hypersensitive sites that could indicate the regulatory region of genes [19]) by employing HINT (i.e., HINT uses hidden Markov models to identify footprints by using strand-specific, nucleosome-sized signals with corrections for ATAC-seq and DNase-seq protocol-specific biases to successfully target CREs) for further processing [20–22]. Our pipeline searches for TF binding sites using TRAP [23] within CREs around accessible genes and calculates an affinity score for each known TF to bind at these particular loci using TEPIC [24,25]. TEPIC uses an exponential decay model that was built under the assumption that regulatory elements close to a gene are more likely important than more distal elements and weighs this relationship accordingly. This allows us to assess TF binding site specific probabilities by using TF binding affinities calculated by TRAP, which uses a biophysical model to assess the strength of the binding energy of a TF to a CREs’ total sequence [23]. Beginning with these CRE candidates, we search for links to possible regulated putative target genes that are differentially expressed between given conditions (e.g., disease and healthy). Approaching the task of linking CREs to target genes, we employ the framework of TEPIC2 [25] and DYNAMITE [25] (feature comparison Suppl. Table 1), which uses a logistic regression model predicting differentially expressed genes across time points and conditions based on TF binding site information to score different TFs according to their contribution to the model and their expression (for a more technical description, see Section Technical Workflow). In general, TF-Prioritizer uses TEPIC and DYNAMITE pairwise of the provided data (i.e., pairwise for each condition and each time point). Based on a background distribution of the scores (combination of differential expression, TEPIC, and DYNAMITE - see Section Discovering Cis-regulatory Elements using a Biophysical Model), TF-Prioritizer computes an empirical p-value reflecting the significance of the results (see Section “An aggregated score to quantify the contribution of a TF to gene regulation”). TF-Prioritizer offers automated access to complementary ChIP-seq data of the prioritized TFs in ChIP-Atlas [17] for validation and shows predicted regulatory regions of target genes using the Integrative Genomics Viewer (IGV) [27–29]. Then, TF-Prioritizer automatically generates a user-friendly and feature-rich web application that could also be used to publish the results as an online interactive report.

**Figure 1:**
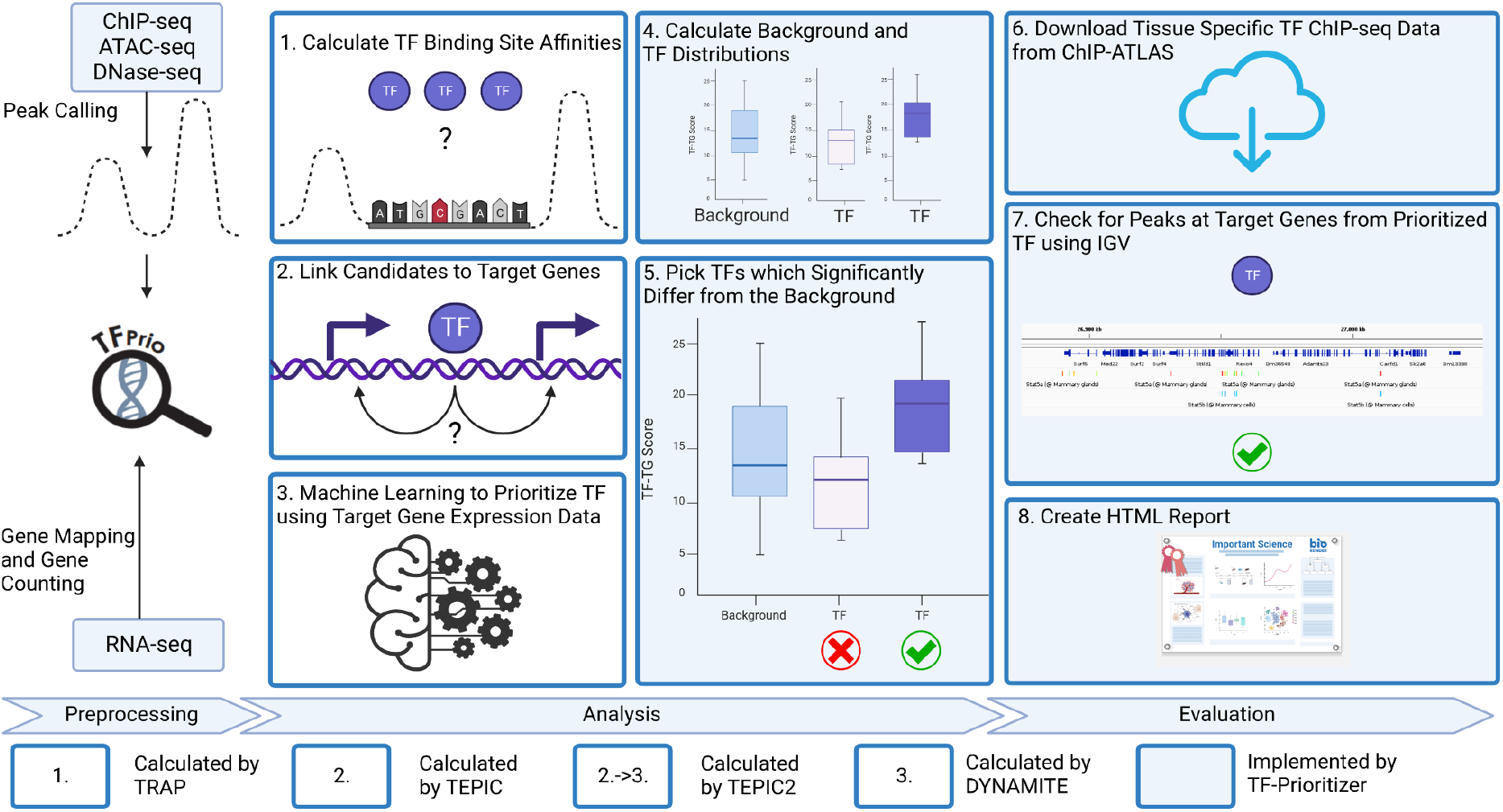
General overview of the TF-Prioritizer pipeline. TF-Prioritizer uses peaks from ChIP-seq or ATAC-seq/DNase-seq and gene counts from RNA-seq. If peaks from the protocols ATAC-seq or DNase-seq were provided, we treat them by using the footprinting method HINT and use the footprints for further processing [20–22]. It then (1) calculates TF binding site affinities using the tool TRAP [23], (2) links candidate regions to potential target genes by employing TEPIC [24], (3) performs machine learning (by using the framework of TEPIC2 [25] and DYNAMITE) to find relationships between TFs and their target genes, (4) calculates background and TF distributions, (5) picks TFs which significantly differ from the background using the Mann-Whitney U test [26] and a comparison between the mean and the median of the background and TF distribution, (6) searches for tissue-specific TF ChIP-seq evaluation data in ChIP-ATLAS [17], (7) creates screenshots using the Integrative Genomics Viewer from predicted regions of interest [27–29], and (8) creates a feature-rich web application for researchers to share and evaluate their results.

To demonstrate the potential and usability of TF-Prioritizer, we use genomic data describing mammary glands in pregnant and lactating mice and compare our analysis to established knowledge [30]. Employing the web application generated by TF-Prioritizer, we found well-studied TFs involved in the mammary gland development process, and we identified additional TFs, which are candidate key factors in mammary gland physiology. Additionally, we use ENCODE cell line data (K562 and MCF-7) to demonstrate the potential and usability of TF-Prioritizer using ATAC-seq, DNase-seq, and HM ChIP-seq data.

## MATERIALS AND METHODS

### Implementation

The main pipeline protocol is implemented in Java version 11.0.14 on a Linux system (Ubuntu 20.04.3 LTS). The pipeline uses subprograms written in Python version 3.8.5, R version 4.1.2, C++ version 9.4.0, and CMAKE version 3.16 or higher. External software that needs to be installed before using TF-Prioritizer can be found on GitHub (see Availability Section). We also provide a bash script “install.sh”, that automatically downloads and installs necessary third-party software and R/Python packages. The web application uses Angular CLI version 14.0.1 and Node.js version 16.10.0. We also provide a dockerized version of the pipeline; it uses Docker version 20.10.12 and Docker-Compose version 1.29.2 (Availability Section). TF-Prioritizer is available as a docker that can be pulled from docker hub and GitHub packages (Availability Section).

### Data processing

#### Mammary gland development and lactation in mice

Data sets (GEO accession id: GSE161620) are processed with the nf-core / RNA-seq [31] and nf-core / ChIP-seq pipelines in their default settings, respectively [32,33]. The FASTQ files of pregnant and lactating mice are processed by Salmon [34] and MACS2 [35] to retrieve raw gene counts and broad peak files.

The dataset spans several time points in mammary gland development from pregnancy to lactation. For each stage, two distinct time points are available: pregnancy day 6 (p6), day 13 (p13), and lactation day 1 (L1), day 10 (L10). For each time point, the dataset contains RNA-seq data and ChIP-seq data for histone modifications H3K27ac and H3K4me3, as well as Pol2 ChIP-seq data (Table 1). We used H3K27ac, H3K4me3, and Pol2 data for creating the model.

**Table 1:** Overview of data sets covering mammary gland development from pregnancy to lactation.

|  | p6 | p13 | L1 | L10 | Sum |
| --- | --- | --- | --- | --- | --- |
| ChIP-seq | 3 | 1 | 8 | 4 | 16 |
| *H3K27ac* |  |  |  |  |  |
| ChIP-seq<br>*H3K4me3* | 2 | 3 | 5 | 0 | 10 |
| ChIP-seq<br>*Pol2* | 2 | 0 | 5 | 4 | 11 |
| RNA-seq | 6 | 8 | 3 | 4 | 21 |

#### ENCODE cell lines

ATAC-seq, DNase-seq, ChIP-seq, and RNA-seq data are downloaded from the ENCODE project for the cell lines K562 (human chronic myelogenous leukemia cell line) and MCF-7 (human breast adenocarcinoma cell line) which are both often used to study cancer biology and have been subjected to a large number of different experimental protocols and assays (Table 2, File identifiers in Suppl. Material 1). (https://www.encodeproject.org/search/?type=Experiment).

**Table 2:**
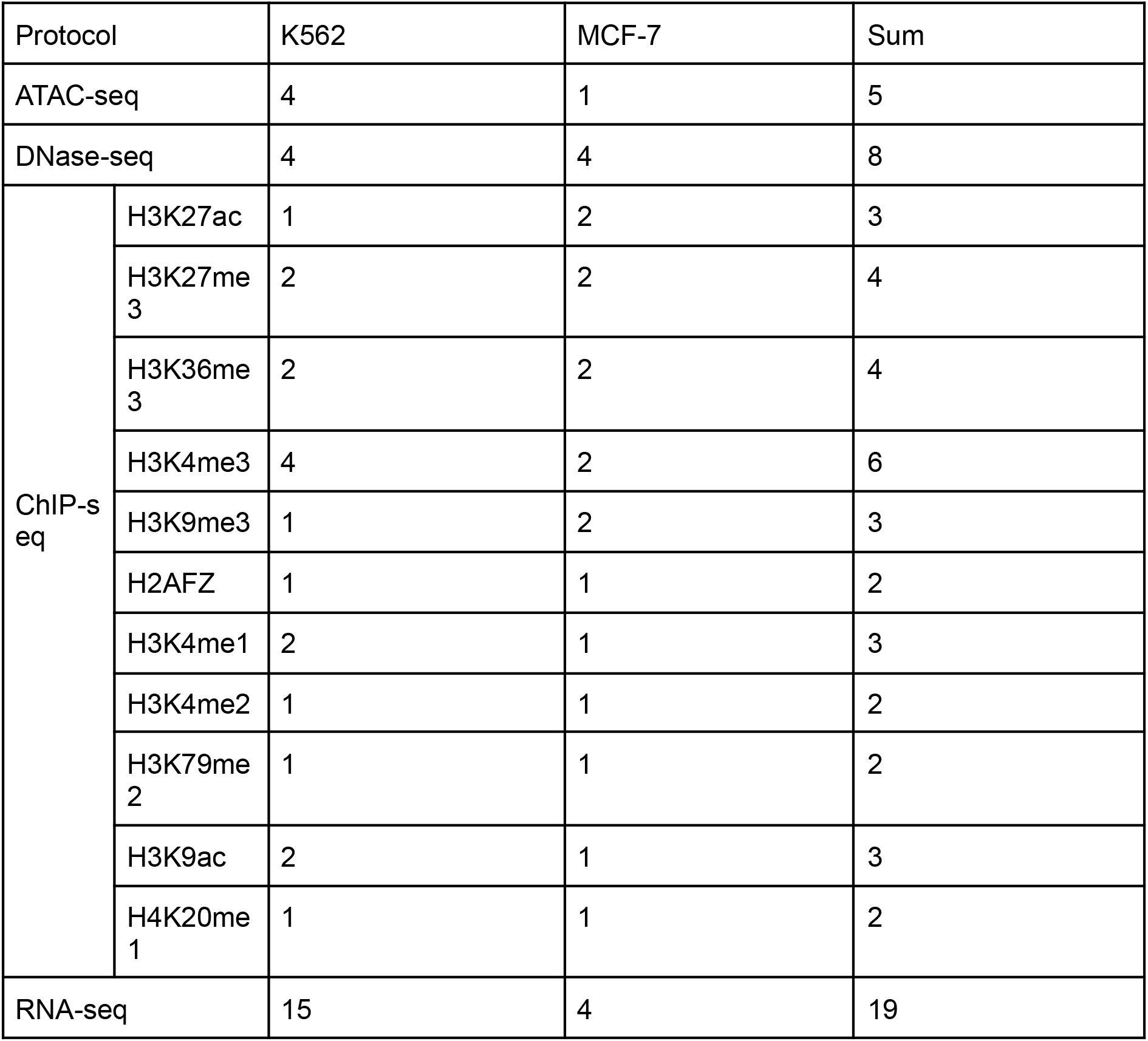
Overview of the data set covering several HM ChIP-seq, ATAC-seq, DNase-seq, and RNA-seq for the cell lines K562 and MCF-7.

| Protocol |  | K562 | MCF-7 | Sum |
| --- | --- | --- | --- | --- |
| ATAC-seq |  | 4 | 1 | 5 |
| DNase-seq |  | 4 | 4 | 8 |
| ChIP-seq | H3K27ac | 1 | 2 | 3 |
|  | H3K27me<sub>3</sub> | 2 | 2 | 4 |
|  | H3K36me<sub>3</sub> | 2 | 2 | 4 |
|  | H3K4me3 | 4 | 2 | 6 |
|  | H3K9me3 | 1 | 2 | 3 |
|  | H2AFZ | 1 | 1 | 2 |
|  | H3K4me1 | 2 | 1 | 3 |
|  | H3K4me2 | 1 | 1 | 2 |
|  | H3K79me<sub>2</sub> | 1 | 1 | 2 |
|  | H3K9ac | 2 | 1 | 3 |
|  | H4K20me<sub>1</sub> | 1 | 1 | 2 |
| RNA-seq |  | 15 | 4 | 19 |

### Technical Workflow

#### Preprocessing

TF-Prioritizer uses peak data from ChIP-seq, ATAC-seq, or DNase-seq and a gene count matrix from RNA-seq as input files (see GitHub repository for detailed formatting instructions). Initially, the pipeline downloads necessary data (gene lengths, gene symbols, and short descriptions of the genes) from BioMart [36]. Optionally, genes with low expression can be removed. TF-Prioritizer uses transcripts per million (TPM) filter of 1 as default to remove TFs that show very low expression and are most probably not relevant. Subsequently, we use DESeq2 to normalize read counts and calculate the log2-fold change (log2fc) [37]. In parallel, TF-Prioritizer preprocesses the peaks by first employing HINT if the provided peak data is labeled as ATAC-seq or DNase-seq to perform footprinting to correct for the biases (i.e., by analyzing chromatin accessibility data in terms of histone modification state, enabling more accurate comparison between the two data types) between the ChIP-seq, ATAC-seq, and DNase-seq protocols [20,38]. TF-Prioritizer then filters blacklisted regions which would likely lead to false positives [39]. Peak files from the same sample group can be merged to significantly reduce the total runtime of the pipeline without affecting the ability of the TF-Prioritizer to identify candidate CREs.

#### Discovering Cis-regulatory Elements using a Biophysical Model

TEPIC links CREs to target genes using a window-based approach (default: 50,000 bp) [24,25] using TRAP, a biophysical model to quantify transcription factor affinity [23]. The window-based approach can be enhanced by providing Hi-C loop data, where the prediction window is extended or limited to a chromatin loop around potential CREs and target genes. TEPIC interprets ChIP-seq signal intensity as a quantitative measure of TF binding strength, which also helps in recovering low-affinity binding sites that would be missed in a classical presence/absence model [24]. The default TEPIC framework searches for dips on top of peaks. However, numerous studies have shown that CREs are often enriched between histone peaks (peak-dip-peak or peak-valley-peak model) [40]. To better accommodate histone modification ChIP-seq data, we thus extended the TEPIC framework to search for transcription factor binding sites (TFBS) between two peaks that have close (default 500 base pairs) genomic positions. TEPIC aggregates individual TF affinities into a TF-Gene score which is the sum of the individual affinities normalized by the length of the considered CREs.

According to the description in Schmidt et al. [41], the TF-Gene score *a*_*w*_(*g,t*) for a gene *g* and a TF *t* in window size *w* is calculated as in Equation 1:

Equation 1: Calculation of the TF-Gene score

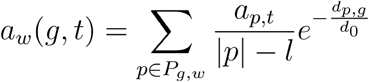

In Equation 1, *a*_*p,t*_ is the affinity of TF *t* in peak *p*. The set of peaks *P*_*g,w*_ contains all open-chromatin peaks in a window of size *w* around the gene *g. d*_*p,g*_ is the distance from the center of the peak *p* to the transcription start site of the gene *g*, and *d*_0_ is a constant fixed at 50,000 bp [42]. The affinities are normalized by peak and motif length, where | *p* | is the length of the peak *p* and *l* is the total length of the motif of TF (see Schmidt et al. for more specific information on how the TF-Gene score is calculated [24,25,41]). Since proximal CREs are expected to have a larger influence on gene expression compared to distal ones, these contributions are weighted following an exponential decay function of genomic distance [25].

We want to point out that the biophysical model calculated by TRAP only returns the center of a potentially large area of high binding energy. The TF is supposed to bind somewhere in this area. In our IGV screenshot, the center of the high binding energy area can appear at a distance up to the window defined by TEPIC. We consider predicted TF peaks as matching if we find TF ChIP-seq peaks inside this window. Following this, we do not expect the predicted TF bindings to overlap exactly with the TF ChIP-seq peaks.

#### An aggregated score to quantify the contribution of a TF to gene regulation

To determine which TFs have a significant contribution to a condition-specific change between two sample groups, we want to consider multiple lines of evidence in an aggregated score. We introduce Transcription Factor Target Gene scores (TF-TG scores, Figure 2) which combine (i) the absolute log2-fold change of differentially expressed genes since genes showing large expression differences are more likely affected through TF regulation than genes showing only minor expression differences; (ii), the TF-Gene scores from TEPIC indicating which TFs likely influence a gene, and (iii) to further quantify this link we also consider the total coefficients of a logistic regression model computed with DYNAMITE [25]. DYNAMITE predicts (high/low) expression of a gene based on the fold changes of TF-Gene scores reported by TEPIC and thus helps to prioritize among multiple potential TFs regulating a gene. We calculate TF-TG scores () for each time point and each type of ChIP-seq data (e.g., different histone modifications) as in Equation 2:

**Figure 2:**
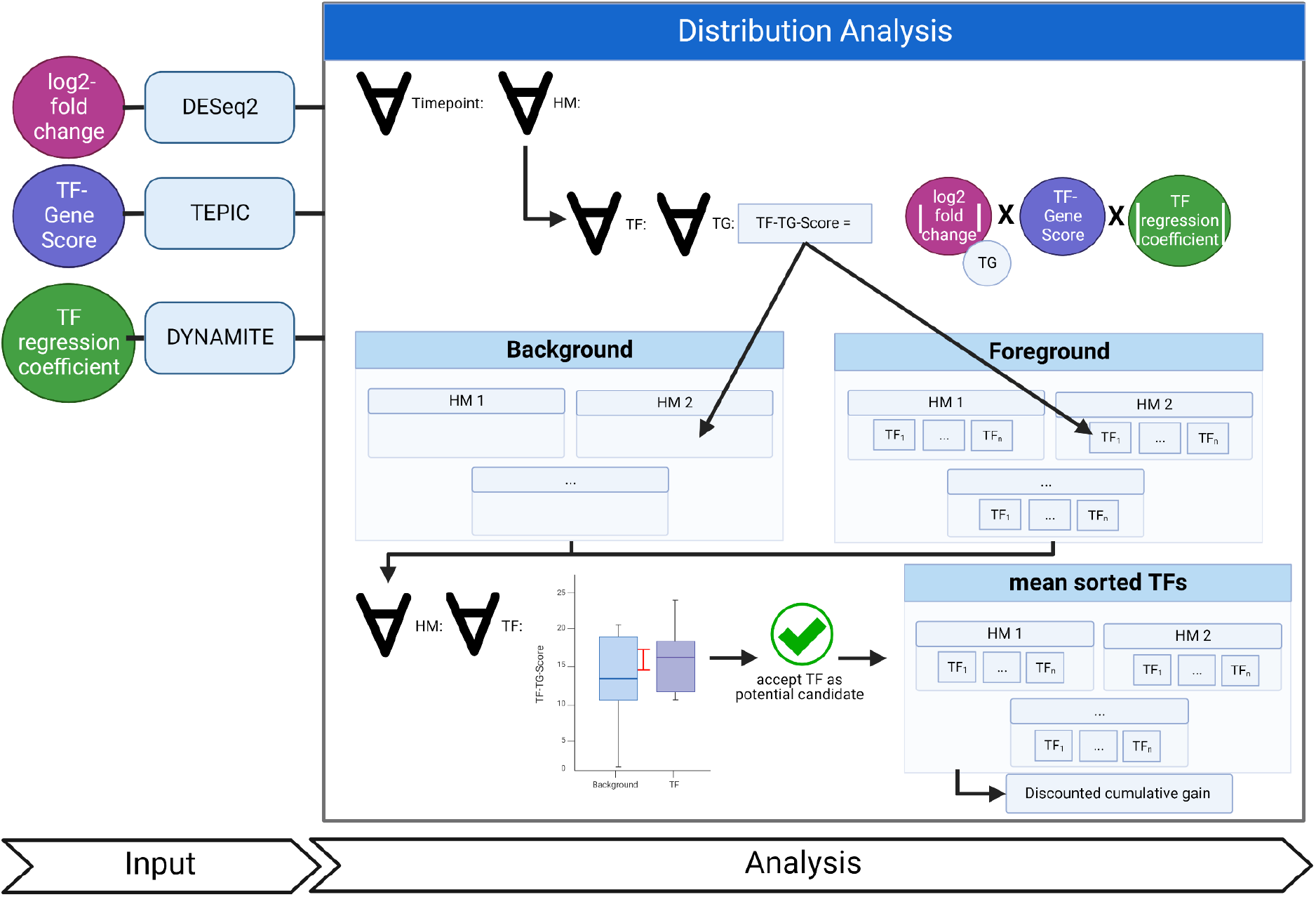
Workflow of the Distribution Analysis to prioritize TFs in a global context by using TF-TG scores. We use several scores conducted by previously performed analysis (see Suppl. Fig. 1), specifically the total log2-fold change (DESeq2), the TF-Gene score (TEPIC), and the total TF regression coefficient (DYNAMITE). We then calculate the TF-TG score for each time point for each TF on each of the TFs predicted target genes (TG) and save it to separate files for the background of each histone modification and for each TF in each histone modification. In the next step, we perform a Mann-Whitney U [43] test between the distribution of the background of the histone modification and the distinct TF distribution of the same histone modification. If the TF passes the Mann-Whitney U test and the median and mean of the TF are higher than the background median and mean, we consider this TF as prioritized for the histone modification. We perform a discounted cumulative gain to receive one list with all prioritized TFs and overall histone modifications.

Equation 2: Calculation of the TF-TG score for each time point and each type of ChIP-seq data :

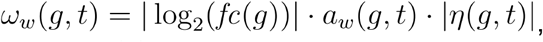

where *fc*(*g*) represents the fold change of the target gene *g* between the two conditions, *a*_*w*_ (*g,t*) the TF-Gene score retrieved by TEPIC as detailed above, *η* (*g,t*) and the total regression coefficient of DYNAMITE’s linear model of the expression of the target gene as a function of the expression of the TF *t*.

#### A random background distribution allows TFprioritzier to exclude spurious results

The ultimate goal of TF-Prioritizer is to identify those TFs that are most likely involved in regulating condition-specific genes. To judge if a specific TF-TG score is meaningful, we generate a background distribution under the hypothesis that the vast majority of TFs will not be condition-specific. Therefore, we generate two different kinds of distributions (see Figure 2): (1) For each HM *m*, a background distribution containing all positive TF-TG scores associated with *m*: *BG*(*m*) = { *ω* _*w*_(*g,t*) | *t* ∈ *TF*(*m*), *g* ∈ *TG*(*t*), *ω* _*w*_(*g,t*) > 0}. Here, *TF*(*m*) denotes the set of TFs that can bind to strands of the DNA modified by *m* and *TG*(*t*) is the set of target genes of the TF *t*. (2) For each HM-TF pair (*m,t*) with *t* ∈*TF*(*m*) a foreground distribution containing all positive TF-TG scores associated with(*m,t*): *FG*(*t,m*) = {*ω*_*w*_(*g,t*) | *g* ∈ *TG*(*t*), *ω* _*w*_ (*g,t*) > 0}. Note that *FG*(*t,m*) ⊆ *BG*(*m*) holds for all HM-TF pairs (*m,t*). We then test each TF distribution of each ChIP-seq against the global distribution matching the ChIP-seq data type. If the p-value of a Mann-Whitney U (MWU) test [43] is below the threshold (default: 0.05) and the median and mean of TF are higher than the background distribution, the TF is recognized as a potential candidate. In the last step, we sort the TFs based on the mean of the TF-TG scores and report the ranks.

We obtain a global list of prioritized TFs across several ChIP-seq data types (e.g., different histone modifications) as follows:

Let *S*(*m*) be the set of transcriptions factors *t* such that the one-sided MWU test between the foreground distribution *FG*(*t,m*) and the background distribution *BG*(*m*) yields a significant *P*-value. For a fixed TF *t* ∈ *S*(*m*), let 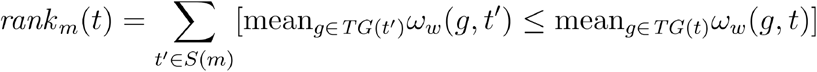 be the rank of *t* in *S*(*m*) w.r.t. the mean TF-TG scores across all target genes, where [·] is the Iverson bracket, i.e., `[true] = 1` and `[false] = 0`. We now compute an overall TF score *f* (*t*) by aggregating the HM-specific ranks as follows:

Equation 3:

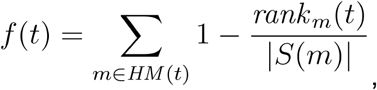

where *HM* (*t*) denotes the set of histone modifications on strands of the DNA where the TF can bind. Note that if *t* ∉*S* (*m*) , rank_*m*_(*t*) is not defined. In this case, we set rank_*m*_(*t*) = |*S* (*m*) | such that the summand for *t* equals 0. Lastly, we sort TFs in ascending order according to the scores *f* (*t*).

#### Discovering each score’s contribution to the global score

To analyze the impact of the different parts of the TF-TG-Score, we permute its components (TF-Score from TEPIC, regression coefficient of DYNAMITE, log2fc of DESeq2). We execute TF-Prioritizer with the exact same configuration but with all possible combinations of the components and compare the prioritized TFs (e.g., solely TF-Score from TEPIC, a combination of TF-Score from TEPIC with the regression coefficient of DYNAMITE, …).

### Validation using independent data from ChIP-Atlas

TF-Prioritizer is able to download and visualize experimental tissue-specific TF ChIP-seq data for prioritized TFs from ChIP-Atlas [17], a public database for ChIP-seq, ATAC-seq, DNase-seq, and Bisulfite-seq data. ChIP-Atlas provides more than 362,121 data sets for six model organisms, i.e., human, mouse, rat, fruit fly, nematode, and budding yeast [44]. TF-Prioritizer automatically visualizes TF ChIP-seq peaks on predicted target sites of prioritized TFs to experimentally validate our predictions. TF-Prioritizer also visualizes experimentally known enhancers and super-enhancers from the manually curated database ENdb [45]. Additionally, experimental data from other databases or experimental data retrieved by own experiments can be supplied and processed by TF-Prioritizer.

By employing TF ChIP-seq data from ChIP-Atlas, TF-Prioritizer is capable of performing a TF co-occurrence analysis of prioritized TFs by systematically comparing the experimentally validated peaks of pairs of prioritized TFs. In a co-occurrence analysis, it is checked what percentage of available peaks of one TF is also found in another TF. TF-Prioritizer returns the percentage of similar peaks between prioritized TFs to discover the co-regulation of TFs. We investigate the co-occurrence of TFs *t*_1_ and *t*_2_ in terms of statistical significance by calculating a log-likelihood score. Let *B* be the set of all TF binding sites and Π (*t*) be the set of peaks for TF *t*. For TF *t*, let *count*(*t*) be the number of binding sites *b* ∈ *B* such that there is a peak π ∈ Π (*t*) within *b*. For a TF-TF pair (*t*_1_,*t*_2_) , let *count*(*t*_1_,*t*_2_) be the number of binding sites *b* ∈ *B* such that there is a peak π_1_ ∈ Π (*t*_1_) and a peak π_2_ ∈ Π (*t*_2_) within *b* then the log-likelihood score *G*^2^ is calculated for the four observations (a) *count*(*t*_1_,*t*_2_) (i.e., *t*_1_ and *t*_2_ are co-occurring), (b) *count*(*t*_1_) − *count*(*t*_1_,*t*_2_) (i.e., *t*_1_ is occurring but *t*_2_ is not), (c) *count*(*t*_2_) − *count*(*t*_1_,*t*_2_) (i.e., *t*_2_ is occurring but *t*_1_ is not), and (d) *count*(*t*_1_,*t*_2_) − *count* (*t*_1_) − *count* (*t*_2_) + | *B* | (i.e., neither *t*_1_ nor *t*_2_ is occurring) with their corresponding expectation values (a) *count* (*t*_1_) · *count* (*t*_2_), (b) *count* (*t*_*1*_) * (| *B* | *count* (*t*_*2*_)), (c) (| *B* | − *count* (*t*_*1*_)) * *count* (*t*_*2*_), and (d) (| *B* | − *count* (*t*_*1*_)) * (| *B* | − *count* (*t*_*2*_) as follows [46–48]:

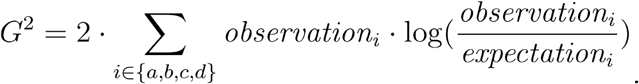

Note that when interpreting each log-likelihood score needs to be brought into relation with the number of peaks found in the respective TFs and also set in relation with the other number of peaks determined in the entire log-likelihood table, as the log-likelihood score may differ from TF-pair to TF-pair. A high log-likelihood score, in combination with a high number of peaks, with respect to the entire log-likelihood table, generally indicates that the co-occurrence relationship is statistically significant and that the two TFs could be functionally related. For further details and explanation of the formula and interpretation, consult [46–48].

#### Explorative analysis of differentially expressed genes

TF-Prioritizer allows users to manually investigate the ChIP-seq signal in the identified CREs of differentially expressed genes. To this end, TF-Prioritizer generates a compendium of screenshots of the top 30 upregulated or downregulated loci (sorted by their total log2-fold change) between two sample groups. Additionally, we allow the user to specify loci that are of special interest (e.g., the CSN family or the *Socs2* locus in lactating mice). TF-Prioritizer then produces screenshots using the TF ChIP-seq data from ChIP-Atlas and visualizes them in a dynamically generated web application. Screenshots are produced using the IGV standalone application [27–29]. TF-Prioritizer also automatically saves the IGV session so the user can further research the shown tracks.

#### Handling missing data

In some cases, not all assay types are available for all samples, or the data does not have the same high quality as the rest of the samples. TF-Prioritizer then skips the grouping of missing data points and can still find meaningful results in the rest of the data. For example, the data for three time points for one histone modification is available, but one time point was missing or discarded. TF-Prioritizer then uses only the three available time points for grouping and downstream processing and analysis.

### Using TF-Prioritizer to investigate gene regulation

We use three approaches to evaluate the biological relevance and statistical certainty of our results: (1) literature research to validate whether the reported TFs are associated with the phenotype of interest, (2) we consider the top 30 target genes with highest affinity values and determine if their expression cluster by condition (note: we do not preselect differentially expressed genes for this analysis but focus on affinities to avoid a circular line of reasoning); we also review the literature and report whether these genes are known to be involved in either pregnancy or mammary gland development/lactation, and (3) validation using independent TF ChIP-seq data from ChIP-Atlas. To conduct the third evaluation, we built region search trees, a balanced binary search tree where the leaves of the tree have a start and end position, and the tree returns all leaves that overlap with a searched region for all chromosomes of the tissue-specific ChIP-Atlas peaks for each available prioritized TF [49]. We then iterate over all predicted regions within the window size defined in TEPIC and determine if we can find any overlapping peaks inside the ChIP-Atlas peaks. If we can find an overlap with a peak defined by the ChIP-Atlas data, we count the predicted peak as a true positive (TP) or else as a false positive (FP). Next, we randomly sample the same number of predicted peaks in random length-matched regions not predicted to be relevant for a TF. If we find an overlap in the experimental ChIP-Atlas data, we consider this region as a false negative (FN) or else as a true negative (TN). Notably, we expect the FN count to be inflated since we considered condition-specific peaks of active CREs. Inactive CREs may very well have TFBS that are not active. Nevertheless, we expect to find more such TFBS in active regions compared to random samples, allowing us to compute sensitivity, specificity, precision, accuracy, and the harmonic mean between precision and sensitivity (F1-score) (see Suppl. Material 2).

### Choice of Parameters

In a pipeline like TF-Prioritizer, the choice of parameters is crucial to retrieve meaningful results. In this section, we explain our choice of parameters. We filter the RNA-seq data by a mean DESeq2 normalized gene count of 50 and a TPM of 1 to exclude noise of very weakly expressed target genes and TFs that are probably not important for the condition but would negatively impact the predictive models. We use the default configurations of TEPIC with the exception of the TF binding site search (i.e., in the histone modification ChIP-seq data, it is important to search for TF binding sites between two peaks that are in close proximity (max. 500 base pairs) to each other (peak-dip-peak or peak-valley-peak model) [40]). The TEPIC2 framework and DYNAMITE were executed in default configurations as provided by the authors. We provide all default parameters in our configuration file.

## RESULTS AND DISCUSSION

We present TF-Prioritizer, which combines data to identify candidate CREs (e.g., ChIP-seq, ATAC-seq, DNase-seq) and RNA-seq to identify condition-specific TF activity. TF-Prioritizer is built on several existing state-of-the-art tools for peak calling, TF-affinity analysis, differential gene expression analysis, and machine learning tools. TF-Prioritizer is the first to jointly consider multiple types of modalities (e.g., different histone marks and/or time series data), provide a joint list of active TFs, and enable the user to see a visualized validation of the predictions in an interactive and feature-rich web application.

### Exploring TFs in mammary tissue during pregnancy and lactation in mice

We used TF-Prioritizer to identify TFs that are known to control mammary gland development and lactation. The tool also identifies TFs that are important in pregnancy, as well as new candidate TFs that have not yet been widely studied. TF-Prioritizer reported 104 TFs, many of which control Rho family GTPase-associated target genes and Casein family genes. TF-Prioritizer was evaluated using experimental TF ChIP-seq data where it showed high sensitivity, specificity, precision, and accuracy (Suppl. Fig. 2, Suppl. Material 2).

#### Prioritized TFs are known to play a role in mammary gland development and lactation

TF-Prioritizer prioritized STAT5, a transcription factor that plays an important role in mammary gland development [30,50,51]. *Stat5* mRNA levels are highly upregulated during the last days of pregnancy and at the beginning of lactation, supporting experimental findings that STAT5 is a key driver of mammary gland development. The predicted target genes of STAT5 show a clear expression separation between pregnancy and lactation (Figure 3 a, b). Peaks were predicted with a sensitivity of 57.8%, specificity of 66.3%, a precision of 78.1%, an accuracy of 60.6%, and an F1 score of 66.5% (Suppl. Fig. 2). Additionally, STAT5 is known to activate the expression of the *Socs2* gene during mammary gland development [52,53]. We can observe predicted peaks of STAT5 near *Socs2*, which could explain the regulation of its expression by STAT5 (Figure 3 c). STAT5 is further known to regulate the expression of the Casein gene family. *Csn2, Csn1s2a, and Csn1s2b* [54] mRNA levels are strongly upregulated during lactation, which could be explained by an activator role of STAT5 at the predicted peaks in their close proximity [55–57] (Figure 3 d, Suppl. Fig. 3, Suppl. Material 3, Sec. STAT5).

**Figure 3:**
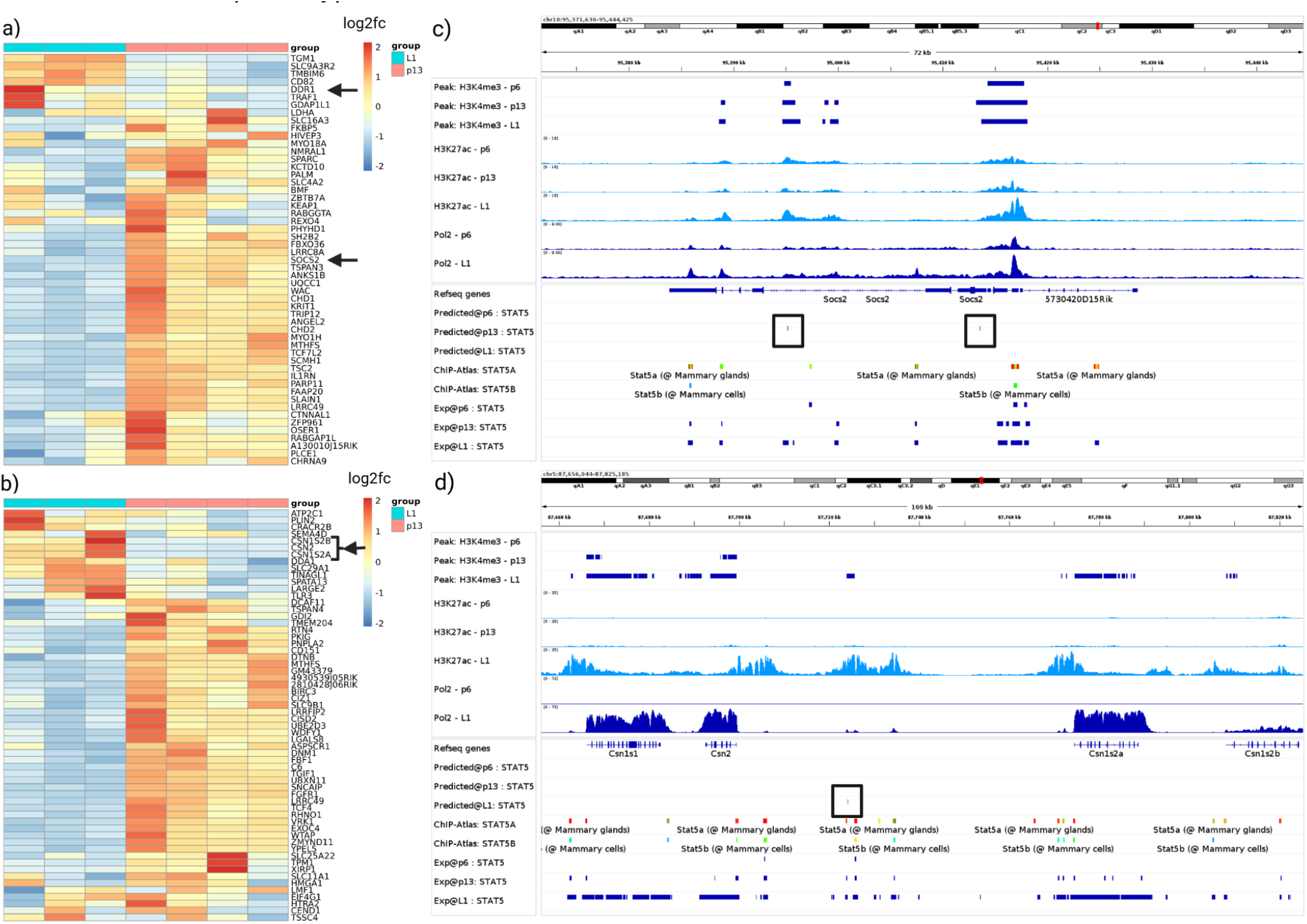
Validation of selected STAT5 target genes. (a) and (b) show heat maps of predicted target genes. We select *Socs2* and *Csn* family genes (black arrows) as they are known to be crucial in either mammary gland development or lactation. In the heatmaps, we can observe a clear separation of these target genes between the time points p13 and L1. Panels (c) and (d) show IGV screenshots of the loci of *Socs2* and the *Csn* family. We included a predicted track in the IGV screenshot that indicates high-affinity binding regions for the TF that are represented by a tick and a black box surrounding it. In (c), we see that we predict peaks in p13 near *Socs2*. From this data, we suggest that *Socs2* mRNA expression is controlled by STAT5 *[52,53]*. In (d), we can observe Pol2 tracks that show a distinct change in the expression of *Csn* family proteins between pregnancy and lactation. This indicates that STAT5 controls the expression of milk proteins.

Additionally, ELF5, another transcription factor that plays an important role in mammary gland development, was predicted to be relevant by TF-Prioritizer. *Elf5* mRNA levels increase at the end of pregnancy and the beginning of lactation, hence supporting ELF5’s role in mammary gland development. Peaks were predicted with a sensitivity of 77.5%, specificity of 80.5%, a precision of 81.6%, an accuracy of 79%, and an F1 score of 79.5% (Suppl. Fig. 2). TF-Prioritizer predicts ELF5 binding sites near *Gli1. Gli1* mRNA levels are downregulated during lactation, and ELF5 is thus probably acting as a suppressor for *Gli1*. Fiaschi et al. showed experimentally that *Gli1*-expressing females were unable to lactate, and milk protein gene expression was essentially absent [58] (Suppl. Figs. 4 and 5, Suppl. Material 3, Sec. ELF5).

TF-Prioritizer further prioritized ESR1 [59] and NFIB *[30]*, both known for their essential function in mammary gland development and lactation (Suppl. Material 3, Sec. ESR1 and NFIB). Our results suggest that the mechanisms of pregnancy, mammary gland development, and lactation could be dependent on Rho GTPase [60,61] and its regulation by several TFs reported here. Experimental validation is needed to elucidate those complex processes further (see Suppl. Material 3, Sec. Rho GTPase’s role in pregnancy, mammary gland development, and lactation) [62].

#### Prioritized novel TFs with a predicted role in pregnancy, mammary gland development, and lactation

We predict two TFs, CREB1 and ARNT, suggesting a role in the processes of pregnancy, mammary gland development, and lactation.

CREB1 binding sites show considerable overlap with binding sites of other TFs known to be involved in mammary gland development and lactation, such as ELF5 (22% of binding sites overlap, log-likelihood score 6,914 with a sample size of 16,531), NFIB (29% binding sites overlap, log-likelihood score 15,793 with a sample size of 23,923), and STAT5A (21% binding sites overlap, log-likelihood score 5,902 with a sample size of 15,180) (see Suppl. Figs. 6. a-c). The co-occurrences could be significant due to the high log-likelihood values with a high sample size in comparison to the whole co-occurrence table. We hypothesize that a correlation of association strength may offer additional evidence for a functional association between TFs. Indeed, CREB1 shows a moderate correlation of binding site affinities with NFIB, STAT5A, STAT5B, and ELF5 (Suppl. Fig 7). Our results suggest that CREB1 regulates a member of the Rho GTPase gene family and a member of the Casein gene family. Since CREB1 has not yet been recognized to contribute to aspects of mammary development and physiology, further experimental validation of our findings is needed (Suppl. Material 3, Sec. CREB1).

Furthermore, the TF ARNT is prioritized along with two cofactors and predicted to be more involved in mammary gland development but less involved in lactation due to its high expression levels during the last state of pregnancy and lower expression during lactation. However, experimental mouse genetics demonstrated that ARNT is not required for mammary development and function [63], suggesting the presence of alternative and compensatory pathways. (Suppl. Material 3, Sec. ARNT).

#### Comparing TF-Prioritizer and diffTF

We compared TF-Prioritizer against the state-of-the-art tool diffTF that prioritizes and classifies TFs into repressors and activators given conditions (e.g., health and disease) [16]. diffTF does not allow multiple conditions or time series data and distinct analysis of histone modification peak data in a single run and does not consider external data for validation. We point out that diffTF cannot use different sample sizes between ChIP-seq and RNA-seq data, i.e., diffTF requires that for each ChIP-seq sample, there is an RNA-seq sample and vice versa. diffTF does not use a biophysical model to predict TFBS but uses general, not tissue-specific, peaks of TF ChIP-seq data and considers all consensus peaks as TFBS [16]. For a comparison of features and technical details, see Suppl. Table 2 and Suppl. Table 3, respectively. Since the diffTF tool does not provide an aggregation approach to different conditions, we aggregate the prioritized TFs the same way as TF-Prioritizer does (i.e., the union of all prioritized TFs overall runs using diffTF’s default q-value cut-off of 0.1) to enhance the comparability of the final results overall conditions. In summary, diffTF prioritized 300 TFs compared to the 104 TFs (including combined TFs like Stat5a..Stat5b that count as one TF in TF-Prioritizer) that TF-Prioritizer reported (Figure 4 a). It thus seems that diffTF is less specific than TF-Prioritizer (see Suppl. Table 4 for a comparison of prioritized TFs). diffTF also finds known TFs that TF-Prioritizer captures (e.g., STAT5A, STAT5B, ELF5, and ESR1) but did not capture the well-known TF NFIB. diffTF also prioritizes CREB1 and ARNT, which in our opinion, are strong candidates for experimental validation. By deploying 20 cores on a general computing cluster, TF-Prioritizer took roughly 7.5 hours to be fully executed, and diffTF took approx. 41 hours to be fully executed. Due to the high number of TFs that are prioritized by diffTF, we ranked the TFs after their p-value (where a low p-value indicates higher evidence that a TF is involved in the processes) provided by diffTF and cut off the exact same amount of TFs (104 TFs) that are prioritized by TF-Prioritizer to make the benchmarking more comparable and interpretable. We observe that the known TFs drop out (e.g., STAT5A, STAT5B, ELF5, NFIB, ESR1) (Figure 4 b). CREB1, which we suggest to be a good candidate for experimental validation, can still be found in diffTFs prediction. Notably, only 22 TFs are prioritized by both TF-Prioritizer and diffTF by using this cutoff.

**Figure 4:**
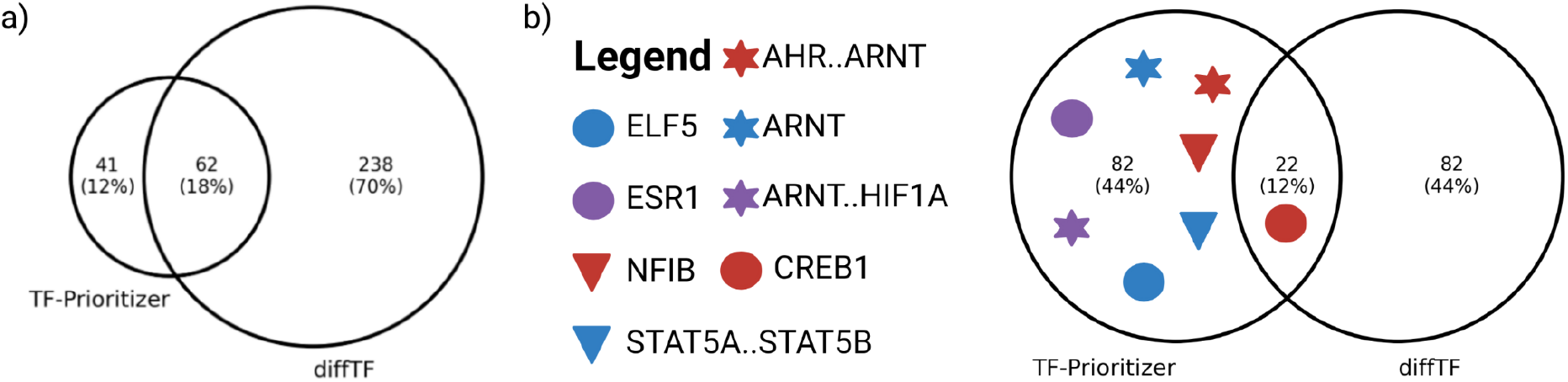
Venn diagram of prioritized TFs by TF-Prioritizer and diffTF. (a) diffTF and TF-Prioritizer found 62 (18.2%) common TFs. diffTF and TF-Prioritizer find known TFs (e.g., STAT5A, STAT5B, ELF5, and ESR1), but diffTF did not capture the well-known TF NFIB. diffTF and TF-Prioritizer both prioritize CREB1 and ARNT as candidates for experimental validation. (b) We ranked the diffTF results by p-value and consider the top 104 (the same amount of TFs that the TF-Prioritizer predicted). Here only CREB1 is still predicted to be important by diffTF - other TFs such as STAT5A..STAT5B, ELF5, and NFIB drop out.

#### Limitations and Considerations

TF-Prioritizer has several limitations. TF-Prioritizer is heavily dependent on the parameters of the state-of-the-art tools it is using, e.g., providing Hi-C data to TEPIC could have a significant impact on the search window while linking potential CREs to target genes. We also point out that we neither have any experimental evidence nor existing literature as proof that the default length of 500 bps of the dip model used in the extended TEPIC framework is the ideal cut-off.

We want to highlight the main disadvantage of using the TF-TG score as we significantly center the surveillance of TF-Prioritizer on genes showing a high fold change or high expression, which does not necessarily mean that those genes are the most relevant for a condition. Also, note that TF binding behavior is regulated by factors we do not observe here, such as phosphorylation. The results of the discounted cumulative gain ranking should be considered with care since the biologically most relevant TFs may manifest in only a subset of ChIP-seq data types.

The calculation of TP, TN, FP, and FN is only an approximation, as to the best of our knowledge, there is no known approach to determine if a CREs or TFBS is active in a condition or not. Sensitivity, specificity, precision, accuracy, and the harmonic mean of precision and sensitivity (F1) differ from TF to TF. We believe this is correlated with the prevalence of the binding sites or the motif specificity. We can also see a decline in the metrics if we look at co-factor regulation (Figure 5. a, AHR..ARNT, ARNT, and ARNT..HIF1A). We experience the highest performance of TF-Prioritizer by looking at TFs where no co-factor regulation is currently known or widely accepted (e.g., CREB1, ELF5, ESR1).

**Figure 5:**
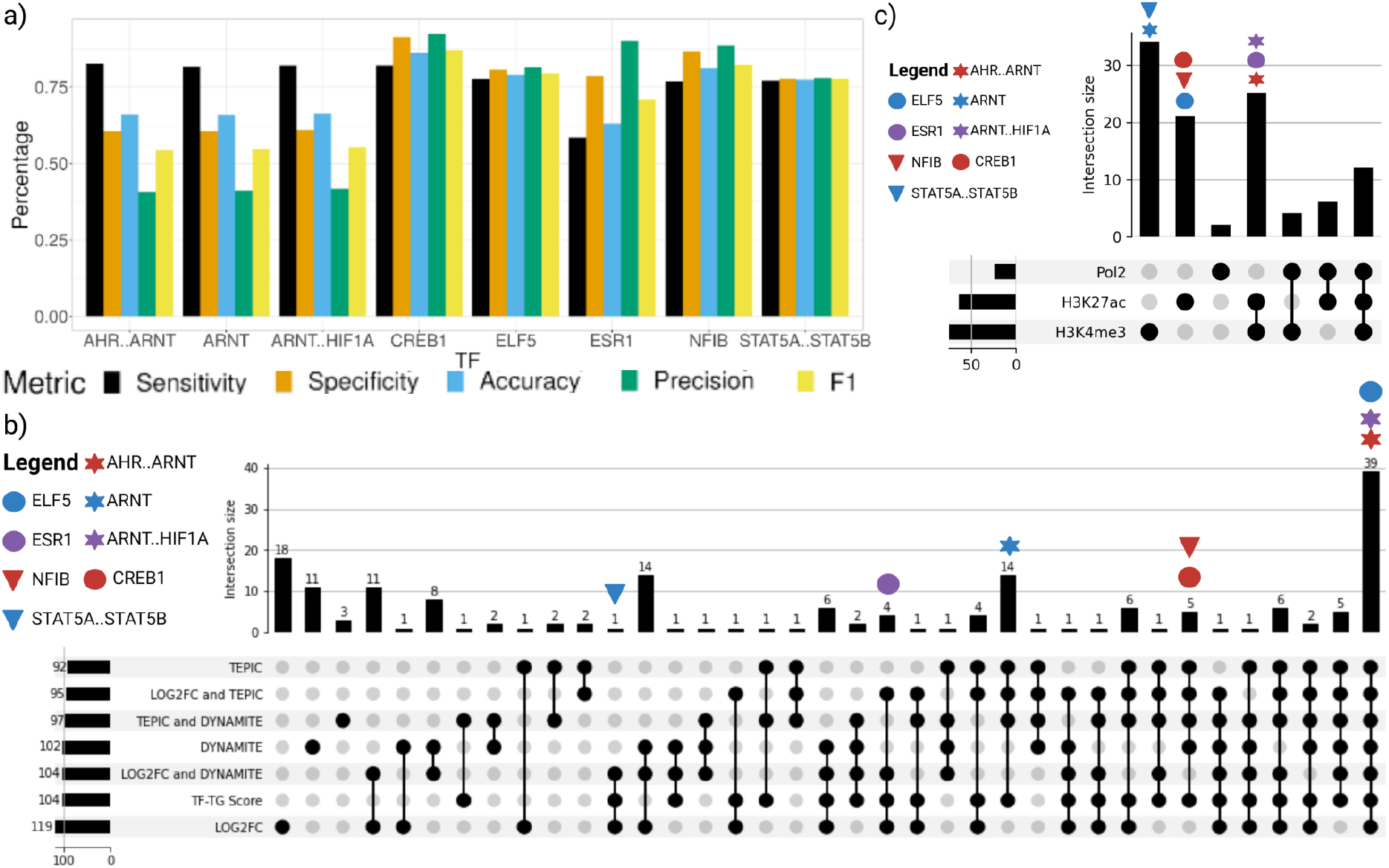
a) Overview of performance metrics of prioritized TFs that were discussed in this manuscript. b) Contributions of individual components of the TF-TG score to the accumulated TF-TG score. We systematically considered different components of the TF-TG score (i.e., the score of TEPIC, LOG2FC, and DYNAMITE) as well as their combinations to determine their importance for the overall results. We find all important TFs exclusively using the TF-TG score. c) Investigation of which TFs are reported in which assay. We can see that the most important TFs only manifest in a subset of HMs.

We further investigated the contribution of every single part of the TF-TG score to the number and quality of the prioritized TFs. To achieve this, we ran every combination of the components of the score (i.e., log2fc, TEPIC, DYNAMITE) with TF-Prioritizer. In Suppl. Table 5 we can see that the distribution analysis filters out about half of the TFs and only returns the most promising TFs. In Figure 5. b, we can see that ELF5, AHR..ARNT, and ARNT..HIF1A manifest in each of the scores independent of any combination. NFIB, CREB1, and ARNT manifest in any score that is related to TEPIC or DYNAMITE. ESR1 manifests in any score that is related to the LOG2FC. STAT5A..STAT5B only manifests in certain combinations of the scores or in the TF-TG score. The LOG2FC alone yields the most prioritized TFs, but at a closer look, the LOG2FC alone would miss NFIB, which is highly relevant in mammary gland development. Looking at this data, we believe that the TF-TG score that combines TEPIC, DYNAMITE, and LOG2FC results in the most promising TFs that are relevant.

In Figure 5. c we can see that STAT5A..STAT5B and ARNT only manifest in the HM H3K4me3. ELF5, CREB1, and NFIB only manifest in H3K27ac. ESR1, AHR…ARNT, and ARNT…HIF1A manifests in both HMs H3K4me3 and H3K27ac. As expected, most TFs only manifest in a subset of HMs, reflecting their association with certain chromatin states [64,65].

### Unraveling the specificity of TFs with respect to HM ChIP-seq, ATAC-seq, and DNase-seq

The ENCODE project generated a plethora of different assays for cell lines such as K562 and MCF-7, which we used here to determine to what extent different protocols (i.e., ATAC-seq, DNase-seq, and HM-ChIP-seq) are suited to reveal condition-specific TFs.

In total, we discovered 381 unique TFs (339 across eleven HM ChIP-seq experiments, 83 in ATAC-seq, and 96 in DNase-seq) if ATAC-seq and DNase-seq open-chromatin peaks were processed with HINT to obtain footprints (Fig. 6, Suppl. Fig. 8 a-c, Suppl. Fig. 9 a-d). Interestingly, the efficacy of footprinting varies between the protocols significantly. Suppl. Fig. 9 shows differences in the number of footprints detected between both protocols. While the number of open chromatin peaks was nearly the same for both protocols, DNase-seq yields fewer footprints compared to ATAC-seq. In general, TF-prioritizer reports more TFs when using footprinting compared to using open chromatin peaks. Many of these overlaps with ChIP-seq TFs, confirming that footprinting is a meaningful strategy(Suppl. Fig. 8 a-b, Fig. 6). We found TFs that can only be detected in a subset of the protocols (Figure 6. a-b, Suppl. Table 6). Using ChIP-seq data, we found the largest number of TFs, likely due to the combination of results from ten different histone modifications and one histone variant, which together cover a wide variety of chromatin states. We found the largest number of detected TFs using the H2AFZ histone variant, possibly due to background peaks because of low antibody sensitivity in this histone variant. Of note, in Suppl. Fig. 10 a and b, we investigated how the number of identified TFs differ when excluding H2AFZ. We can see a decrease in the total number of prioritized TFs in ChIP-seq from 339 to 301. We further examined how the number of identified TFs changes when only employing frequently studied HM ChIP-seq data from H3K27ac, H3K4me1, and H3K4me3 (Suppl. Fig. 10 c and d). We can observe a decrease in identified TFs from 339 to 152 but, again, the overlap with ATAC-seq and/or DNase-seq drops. H2AFZ is predominantly found in CREs and is also associated with cancer [66]. Since we have only investigated cancer cell lines, it remains unclear if this histone variant is generally highly informative of TF binding or if this is limited to cancer cells. Surprisingly, DNase-seq and ATAC-seq show a comparably small overlap even though both protocols are aimed at measuring chromatin accessibility. This corroborates earlier findings where it was observed that both protocols reveal assay-specific sites that contribute to predicting gene expression [67].

**Figure 6:**
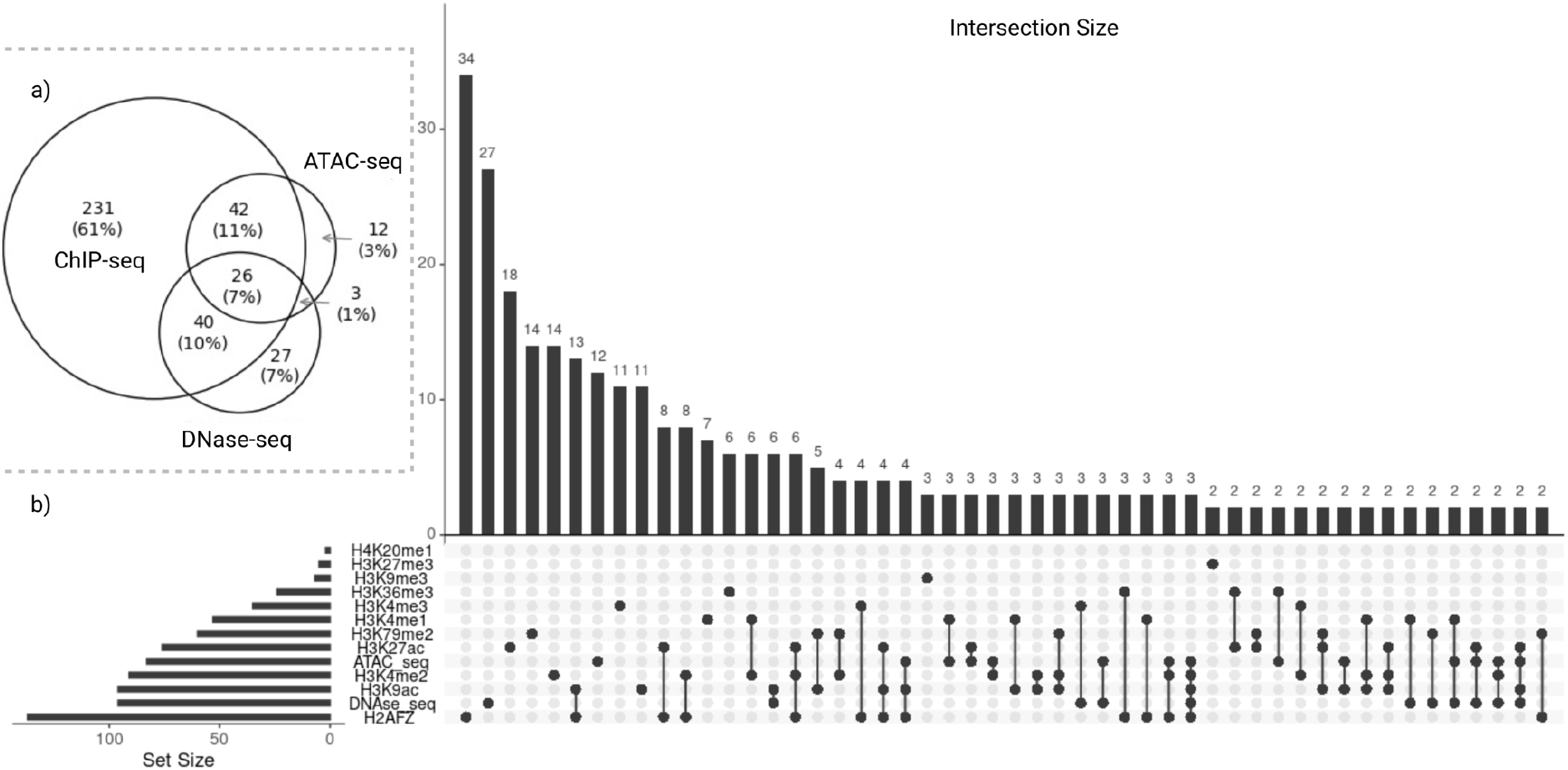
Guide to determine which experiments fit best by the usage of ATAC-seq, DNase-seq, or several histone modifications. a) We combined all HM ChIP-seq data and investigated the overlap with ATAC-seq and DNase-seq. We found that ATAC-seq and ChIP-seq have a bigger overlap than ATAC-seq and DNase-seq. We found 26 TFs that are prioritized by all three protocols. b) We separated the TFs of the HM ChIP-seq data in which HM they were discovered. We can see huge differences between the HMs (e.g., while we can discover 137 TFs in H2AFZ, we can only discover two in H4K20me1).

Indeed, some TFs known to be important for both cancer cell lines were reported through several protocols, while others were reported by only one protocol. For instance, we found MYC, a key TF for cell proliferation in K562 and MCF-7 cells [68,69], was highly ranked in ATAC-seq and HM ChIP-seq (H3K4me2, H3K79me2). Conversely, GATA1, another TF important for cell differentiation in K562 [70,71], was prioritized only by DNase-seq. GATA1 regulates MYB, a key haematopoietic TF involved in stem cell self-renewal and lineage decisions that was prioritized in HM ChIP-seq (H2AFZ, H3K27ac, H3K4me2) [71,72]. TF-Prioritizer found many members of the SP (SP1, SP2, SP3, SP4, SP8, and SP9) and KLF (KLF1, KLF2, KLF3, KLF4, KLF6, KLF7, KL8, KLF9, KLF10, KLF11, KLF12, KLF14, KLF15, and KLF16) family to be important for K562 cell differentiation in a plethora of HM ChIP-seq, ATAC-seq, and DNase-seq experiments. Notably, TF-Prioritizer uses an individual TF energy pattern during the calculation of TF affinity to potential binding (i.e., TRAP) for each TF of a TF family. The incorporation of TF expression data in our score further boosts this differentiation between TFs of the same family. We identified six out of 9 TFs from the SP TF family and 14 out of 16 TFs from the KLF TF family [73]. Hu et al. found that the SP and KLF TF families are most important in erythroid differentiation in K562 cells [74] and that SP1 and SP3 are involved in activating GATA1 [75].

We further investigated if TF-Prioritizer found biologically relevant TFs for the MCF-7 cell line. We found ELF5, an important TF in breast cancer, to be prioritized in ATAC-seq, DNase-seq, and HM ChIP-seq (H2AFZ). This is of particular interest, as ELF5 is a strong biomarker in breast cancer, and TF-Prioritizer is capable of prioritizing ELF5 in the ATAC-seq, DNase-seq, and ChIP-seq [76–78]. Piggin et al. also postulated that ELF5 modulates the estrogen receptor [78]. TF-Prioritizer found certain estrogen receptors (e.g., ESR2, ESRRG) to be relevant for cell differentiation in MCF-7. Estrogen receptor proteins are highly relevant in breast cancer [79,80]. The TF GATA3 was also predicted (ATAC-seq, H3K27ac, H3K9ac) to be important for cell differentiation in MCF-7. GATA3 is a key player when it comes to cell differentiation in the MCF-7 cell line [81,82] and a regulator of estrogen receptor proteins [83]. FOXA1, predicted by TF-Prioritizer (ATAC-seq), is important in cell differentiation for MCF-7 cell lines, is a critical determinant of estrogen receptor function, and affects the proliferation activity of breast cancer [84,85].

## CONCLUSION AND OUTLOOK

TF-Prioritizer is a pipeline that combines RNA-seq and ChIP-seq data to identify condition-specific TF activity. It builds on several existing state-of-the-art tools for peak calling, TF-affinity analysis, differential gene expression analysis, and machine learning tools. TF-Prioritizer is the first tool to jointly consider multiple types of modalities (e.g., different histone marks and/or time series data) and provide a summarized list of active TFs. A particular strength of TF-Prioritizer is its ability to integrate all of this in an automated pipeline that produces a feature-rich and user-friendly web report. It allows interpreting results in the light of experimental evidence (TF ChIP-seq data) either retrieved automatically from ChIP-Atlas or user-provided and processed into genome browser screenshots illustrating all relevant information for the target genes. Our approach was heavily inspired by DYNAMITE [25,86], which follows the same goal but requires manually performing all necessary steps.

We show that TF-Prioritizer is capable of identifying already known and validated TFs (e.g., STAT5, ELF5, NFIB, ESR1) that are involved in the process of mammary gland development or lactation and their experimentally validated target genes (e.g., *Socs2, Csn* milk protein family, Rho GTPase associated proteins). Furthermore, we prioritized some not yet recognized TFs (e.g., CREB1, ARNT) that we suggest as potential candidates for further experimental validation. These results led us to hypothesize that the Rho GTPases undergo major changes in their tasks during the stages of pregnancy, mammary gland development, and lactation, which are regulated by TFs.

In conclusion, each protocol and histone modification can unravel unique transcription factor binding sites that provide insight into gene regulatory mechanisms. It is our opinion that employing TF-Prioritizer on as many protocols and HM ChIP-seq experiments as possible could improve our understanding of given conditions.

In the future, we plan to extend TF-Prioritizer to more closely explore the combined effects of enhancers, which are often non-additive, as suggested by our current model [87]. We further plan to test the functionality of TF-Prioritizer on ATAC-seq data and to apply TF-Prioritizer in a single-cell context where histone ChIP-seq is currently hard to retrieve. Furthermore, we plan to include a more detailed ranking of the prioritized TFs. We plan to offer the user the ability to apply raw FASTQ files to TF-Prioritizer, where quality checks of the data will be performed. In summary, TF-Prioritizer is a powerful functional genomics tool that allows biomedical researchers to integrate large-scale ChIP-seq and RNA-seq data, prioritize TFs likely involved in condition-specific gene regulation, and interactively explore the evidence for the generated hypotheses in the light of independent data.

## Supporting information

Supplementary Data is available at biorxiv online.

## AVAILABILITY AND REQUIREMENTS

The source code of the pipeline is freely available at GitHub: https://github.com/biomedbigdata/TF-Prioritizer

The report on the pregnant and lactating mice data set is available at: https://exbio.wzw.tum.de/tfprio/mouse/#/

Mouse pregnancy and lactation data: https://www.ncbi.nlm.nih.gov/geo/query/acc.cgi?acc=GSE161620

Mouse TF ChIP-seq data on STAT: https://www.ncbi.nlm.nih.gov/geo/query/acc.cgi?acc=GSE82275 https://www.ncbi.nlm.nih.gov/geo/query/acc.cgi?acc=GSE84115 https://www.ncbi.nlm.nih.gov/geo/query/acc.cgi?acc=GSE37646

ChIP-seq, ATAC-seq, and DNase-seq data from K562 and MCF-7 cell lines (Suppl. Material 1 for file identifiers): https://www.encodeproject.org/search/?type=Experiment

The report on the ATAC-seq, DNase-seq, and ChIP-seq is available at:

ATAC-seq: https://exbio.wzw.tum.de/tfprio/cancer/atac/#/

DNase-seq: https://exbio.wzw.tum.de/tfprio/cancer/dnase/#/

ChIP-seq: https://exbio.wzw.tum.de/tfprio/cancer/chip/#/

Table to determine which HM ChIP-seq, ATAC-seq, and DNase-seq data can be used to unravel which TFs: https://figshare.com/articles/dataset/protocols_hms_to_unraveledTFs_tsv/21941213/2

Docker images:

GitHub packages (only accessible via GitHub command line) https://raw.githubusercontent.com/biomedbigdata/TF-Prioritizer/pipeJar/docker.py

docker hub https://hub.docker.com/r/nicotru/tf-prioritizer

Pipeline registrations:

bio.tools https://bio.tools/tf-prioritizer

SciCrunch.org RRID:SCR_023222

workflowhub.eu https://workflowhub.eu/workflows/433 Project name: TF-Prioritizer

Project home page: https://github.com/biomedbigdata/TF-Prioritizer

Operating system(s): Linux

Programming language: Java

Other requirements: Java version 11.0.14 or higher, Python version 3.8.5 or higher, R version 4.1.2 or higher, C++ version 9.4.0 or higher, CMAKE version 3.16 or higher, Angular CLI version 14.0.1 or higher, Node.js version 16.10.0 or higher, Docker version 20.10.12 or higher, and Docker-Compose version 1.29.2 or higher.

Open source license: GNU GPL v. 3.0

## SUPPLEMENTARY DATA

Supplementary Data is available at GigaScience online. https://docs.google.com/document/d/18ErBhbZ9IW6SLeRTF2AB7PF7UFJu6KcQ48Z4JcZD4Xk/edit?usp=sharing

## ABBREVIATIONS

Abbreviation: **Description**
Ahr: Aryl Hydrocarbon Receptor
Arhgap12: Rho GTPase Activating Protein 12
Arhgap39: Rho GTPase Activating Protein 39
Arhgap9: Rho GTPase Activating Protein 9
Arhgef1: Rho Guanine Nucleotide Exchange Factor 1
Arhgef18: Rho/Rac Guanine Nucleotide Exchange Factor 18
Arhgef2: Rho/Rac Guanine Nucleotide Exchange Factor 2
Arhgef40: Rho Guanine Nucleotide Exchange Factor 40
Arhgef9: Cdc42 Guanine Nucleotide Exchange Factor 9
Arnt: Aryl Hydrocarbon Receptor Nuclear Translocator
CREs: cis-regulatory elements
Creb1: CAMP Responsive Element Binding Protein 1
Csn: Casein proteins
Csn1s2a: Casein Alpha S2 Like A
Csn1s2b: Casein Alpha S2 Like B
Csn2: Casein Beta
Csnk1e: Casein Kinase 1 Epsilon
Csnk2a2: Casein Kinase 2 Alpha 2
Csnk2b: Casein Kinase 2 Beta
Ddr1: Discoidin Domain Receptor Tyrosine Kinase 1
Elf5: E74 Like ETS Transcription Factor 5
Esr1: Estrogen Receptor 1
Ets2: ETS Proto-Oncogene 2, Transcription Factor
F1-score: harmonic mean between precision and sensitivity
FN: false negatives
FP: false positives
Gli1: GLI Family Zinc Finger 1
HM: histone modification
Hif1a: Hypoxia Inducible Factor 1 Subunit Alpha
IGV: Integrative Genome Viewer
Igfals: Insulin Like Growth Factor Binding Protein
Acid: Labile Subunit
L1: lactation day 1
L10: lactation day 10
Lcp1: Lymphocyte Cytosolic Protein 1
MWU: Mann-WhitneyU test
Nfib: Nuclear Factor I B
Socs2: Suppressor Of Cytokine Signaling 2
Stat5 (composition of Stat5a and Stat5b): Signal Transducer And Activator Of Transcription 5A + Signal Transducer And Activator Of Transcription 5B
TF: transcription factor
TF-Gene: score retrieved by TEPIC
TF-TG: score retrieved by the Distribution Analysis
TFBS: transcription factor binding sites
TG: target gene
TP: true positives
TPM: transcripts per million
Tp53: Tumor Protein P53
log2fc: log2 fold-change
p13: pregnancy day 13
p6: pregnancy day 6

## AUTHORS’ CONTRIBUTION

MH, NT, FS, JB, MS, DB, LH, and ML drafted the concept for this pipeline. MH, NT, OL, and KY implemented the pipeline. MH, LS, and NT conceptualized and implemented the ATAC-seq and DNase-seq integration. JJ and HKL created the experimental data in the laboratory. JJ, HKL, LLW, KY, and NB prepared the data and performed quality checks. MH, NT, DB, LH, and ML wrote the manuscript. All authors reviewed the manuscript and approved it.

## ACKNOWLEDGMENTS

We want to thank Christina Trummer for designing the TF-Prioritizer logo. We want to thank Andreas Niekler for his help with the statistical analysis of the co-occurrence analysis.

Figures were created with https://www.biorender.com. Parts of the images were designed using resources from Flaticon.com.

