## Supplementary Data is available at biorxiv online. for "TF-Prioritizer: a java pipeline to prioritize condition-specific transcription factors"

### *Supplementary material for:* TF-Prioritizer: a java pipeline to prioritize condition-specific transcription factors

[*]The authors wish to be known that in their opinion the first two authors should be considered as shared first authors

[†]The authors wish to be known that in their opinion the last two authors should be considered as shared last authors

*** contact:*

### ABSTRACT

#### Background

**Eukaryotic gene expression is controlled by cis-regulatory elements (CREs), including promoters and enhancers, which are bound by transcription factors (TFs). Differential expression of TFs and their binding affinity at putative CREs determine tissue- and developmental-specific transcriptional activity. Consolidating genomic data sets can offer further insights into the accessibility of CREs, TF activity, and, thus, gene regulation. However, the integration and analysis of multi-modal data sets are hampered by considerable technical challenges. While methods for highlighting differential TF activity from combined chromatin state data (e.g., ChIP-seq, ATAC-seq, or DNase-seq) and RNA-seq data exist, they do not offer convenient usability, have limited support for large-scale data processing, and provide only minimal functionality for visually interpreting results.**

#### Results

**We developed TF-Prioritizer, an automated pipeline that prioritizes condition-specific TFs from multi-modal data and generates an interactive web report. We demonstrated its potential by identifying known TFs along with their target genes, as well as previously unreported TFs active in lactating mouse mammary glands. Additionally, we studied a variety of ENCODE data sets for cell lines K562 and MCF-7, including twelve histone modification ChIP-seq as well as ATAC-seq and DNase-seq datasets, where we observe and discuss assay-specific differences.**

#### Conclusion

**TF-Prioritizer accepts ATAC-seq, DNase-seq, or ChIP-seq and RNA-seq data as input and identifies TFs with differential activity, thus offering an understanding of genome-wide gene regulation, potential pathogenesis, and therapeutic targets in biomedical research.**

#### Supplementary Figure 1: Technical Workflow

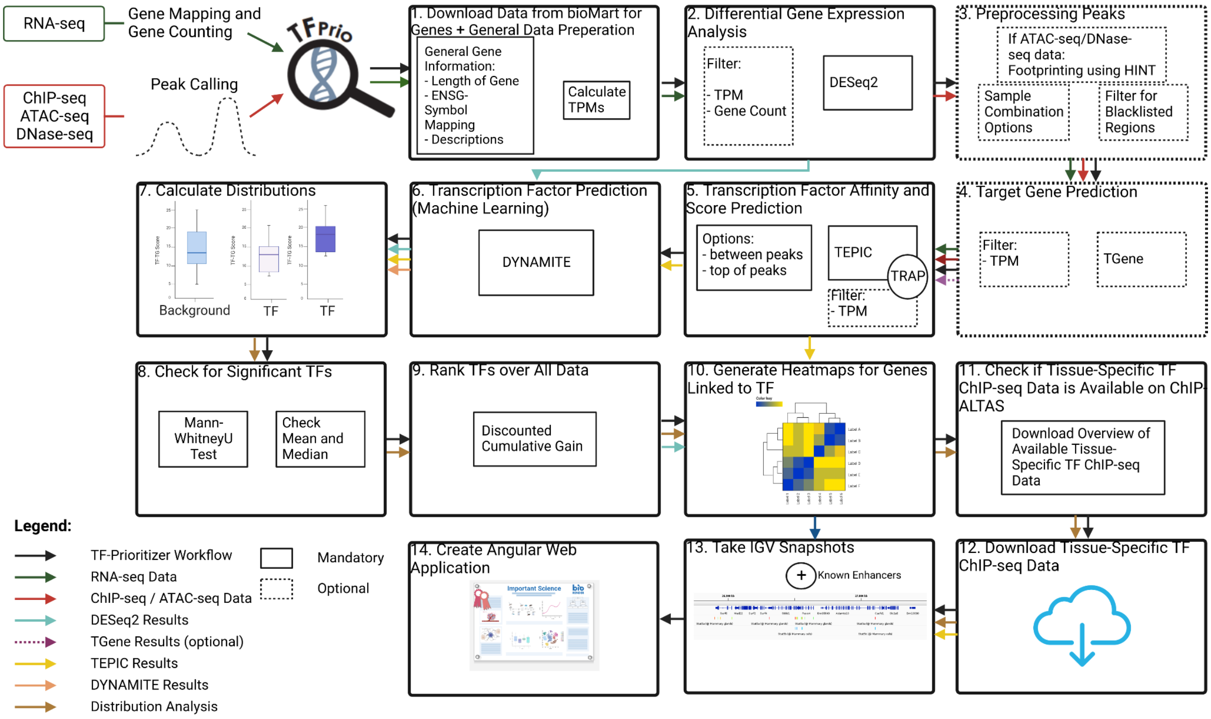

**Supplementary Figure 1:** TF-Prioritizer uses nf-core ChIP-seq / ATAC-seq and nf-core RNA-seq preprocessed data as input files (see GitHub repository for detailed formatting instructions). More specifically, broad peaks and gene counts. (1) Once started, the pipeline downloads necessary data (gene lengths, gene symbols, and short descriptions of the genes) from bioMart [1]. (2) The user can then decide to use a transcript per million (TPM) filter or a gene count filter to filter before DESeq2 usage. We also allow for batch correction in DESeq2. TF-Prioritizer uses a TPM filter of 1 as default. DESeq2 normalizes and calculates the log2 fold change (log2fc) from raw gene count data [2]. If the user used ATAC-seq as an input, we use the footprint method HINT [3–5] to process the peaks, for this process we additionally expect BAM files in the same directory format as the peaks from the user. In parallel, (3) TF-Prioritizer preprocesses the ChIP-seq broad peaks by filtering blacklisted regions [6]. We recommend using the sample combination option to combine similar broad peak samples into one peak file, as the total runtime of the pipeline is reduced significantly without losing the quality of the data. (4) Optionally, the user can decide to use TGene to predict links between target genes and regulatory elements combining distance and histone/expression correlation [7]. If the TGene option is not activated, TEPIC, executed in the next step of the pipeline, uses a window-based approach to link regulatory elements to target genes [8,9]. (5) TEPIC uses TRAP, an approach that quantifies transcription factor affinity scores based on a biophysical model for regulatory regions [10]. TEPIC "computes TF affinities and uses open-chromatin/HM signal intensity as quantitative measures of TF binding strength". TEPIC uses "machine learning to find low-affinity binding sites to improve the ability to explain gene expression variability compared to the standard presence/absence classification of binding sites" [8]. In addition, especially for histone modification ChIP-seq data, we extended the TEPIC framework so that it can also search for transcription factor binding sites (TFBS) between two peaks that have close (~500 bps) genomic positions (default: search between two peaks). (6) The pipeline then executes DYNAMITE an approach that uses a "sparse logistic regression classifier to infer TFs related to gene expression changes between samples" [9]. (7) We added a distribution analysis to the pipeline to further prioritize TFs depending on their distribution compared to the global distribution using (8) a Mann-Whitney U test [11] and the comparison of the means and the medians (for details see Materials and Methods Distribution Analysis Section). (9) We then use a discounted cumulative gain approach to retrieve a global ranking (overall histone modification data provided) of prioritized TFs (see Materials and Methods Discounted Cumulative Gain Section). (10) In the following, TF-Prioritizer generates condition-specific and histone-modification-specific heatmaps for prioritized TFs and their predicted target genes. (11) We then check if we can find publicly available tissue-specific TF ChIP-seq data from ChIP-ATLAS [12] and (12) download the files. (13) Afterward, we take screenshots using the IGV [13–15]. (14) In the last step, we conclude all analysis and plots in form of an easy-to-use HTML report that could also be used as a webpage.

#### Supplementary Figure 2: Confusion matrices and statistical measures

a) STAT5A..STAT5B

| | Experimental | |
|---|---|---|
| Predicted | True | False |
| True | 54,805 | 15,319 |
| False | 39,866 | 30,258 |

| | | | |
|---|---|---|---|
| Sensitivity | 57.89% | Specificity | 66.39% |
| Precision | 78.15% | Accuracy | 60.65% |
| F1-Score | 66.51% | | |

b) ELF5

| | Experimental | |
|---|---|---|
| Predicted | True | False |
| True | 29,864 | 6,737 |
| False | 8,636 | 27,965 |

| | | | |
|---|---|---|---|
| Sensitivity | 77.57% | Specificity | 80.59% |
| Precision | 81.59% | Accuracy | 79.00% |
| F1-Score | 79.53% | | |

c) ESR1

| | Experimental | |
|---|---|---|
| Predicted | True | False |
| True | 9,214 | 1,003 |
| False | 6,459 | 3,758 |

| | | | |
|---|---|---|---|
| Sensitivity | 58.79% | Specificity | 78.93% |
| Precision | 90.18% | Accuracy | 63.48% |
| F1-Score | 71.18% | | |

d) NFIB

| | Experimental | |
|---|---|---|
| Predicted | True | False |
| True | 104,007 | 13,330 |
| False | 30,954 | 86,383 |

| | | | |
|---|---|---|---|
| Sensitivity | 77.06% | Specificity | 86.63% |
| Precision | 88.64% | Accuracy | 81.13% |
| F1-Score | 82.45% | | |

e) CREB1

| | Experimental | |
|---|---|---|
| Predicted | True | False |
| True | 45,263 | 3,699 |
| False | 9,895 | 39,067 |

| | | | |
|---|---|---|---|
| Sensitivity | 82.06% | Specificity | 91.35% |
| Precision | 92.45% | Accuracy | 86.12% |
| F1-Score | 86.94% | | |

f) ARNT

| | Experimental | |
|---|---|---|
| Predicted | True | False |
| True | 17,083 | 24,403 |
| False | 3,811 | 37,675 |

| | | | |
|---|---|---|---|
| Sensitivity | 81.76% | Specificity | 60.69% |
| Precision | 41.18% | Accuracy | 66.00% |
| F1-Score | 54.77% | | |

e) ARNT..HIF1A

| | Experimental | |
|---|---|---|
| Predicted | True | False |
| True | 59,919 | 83,115 |
| False | 45,076 | 97,958 |

| | | | |
|---|---|---|---|
| Sensitivity | 57.07% | Specificity | 54.10% |
| Precision | 41.89% | Accuracy | 55.19% |
| F1-Score | 48.32% | | |

g) AHR..ARNT

| | Experimental | |
|---|---|---|
| Predicted | True | False |
| True | 65,503 | 94,477 |
| False | 38,243 | 121,737 |

| | | | |
|---|---|---|---|
| Sensitivity | 63.14% | Specificity | 56.30% |
| Precision | 40.94% | Accuracy | 58.52% |
| F1-Score | 49.68% | | |

**Supplementary Figure 2:** We showcase the discussed TFs and their statistical metrics. We can see the confusion matrix for each TF. We also provide sensitivity, specificity, precision, accuracy, and F1-score. We can see that the metrics differ vastly between the TFs. There is a drop in all metrics when it comes to co-factors. We believe that more research is necessary to obtain better predictions for co-factor TFs.

#### Supplementary Figure 3: IGV screenshot of STAT5 target gene DDR1

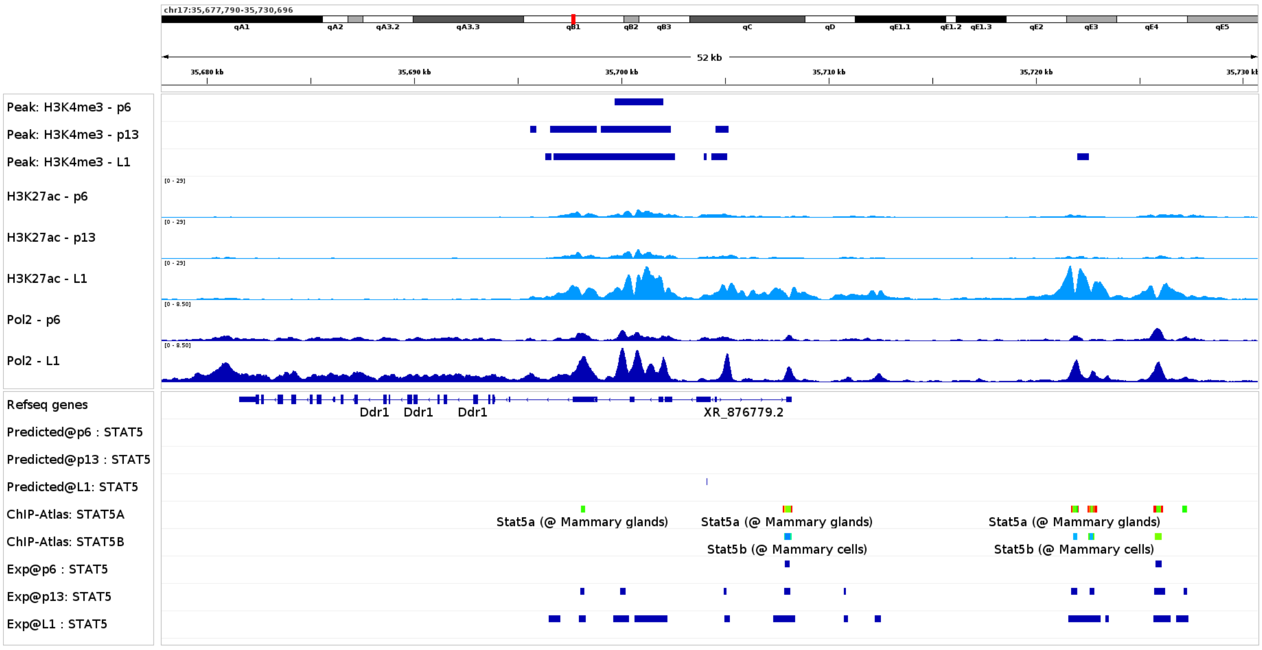

**Supplementary Figure 3:** We show the predicted peaks and experimental signals of DDR1. We can see higher Pol2 signals in the area of DDR1 due to higher expression during lactation. We furthermore can also observe a predicted peak of STAT5 in the DDR1 region. Past research showed that DDR1 is necessary to maintain STAT5 signaling during lactation.

#### Supplementary Figure 4: Heatmaps and target genes of ELF5

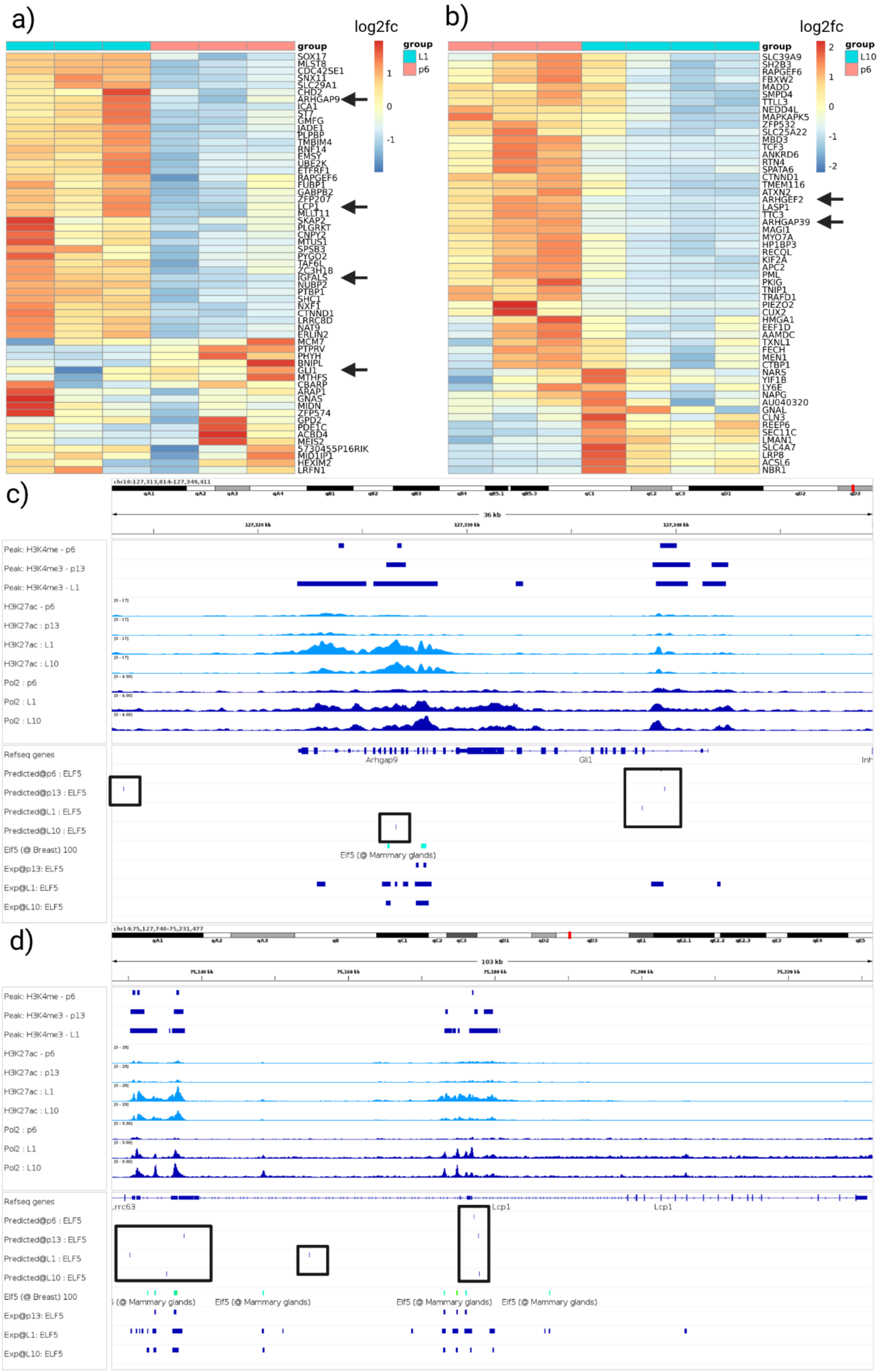

**Supplementary Figure 4:** Validation of selected target genes for *Elf5*. (a) and (b) show heat maps of predicted target genes. We select *Gli1*, *Lcp1*, and *Igfals* (black arrows) as they are already known to be crucial in either mammary gland development or lactation. We further select the genes *Arhgap9*, *Arhgef2*, and *Arhgap39* (black arrows) that are known to be essential for Rho GTPases due to their studied role in epithelial morphogenesis during mammary gland development [54,55]. In the heatmaps, we can observe a clear separation of these target genes between the time points p6-L1 and p6-L10. (c) and (d) show IGV screenshots of *Arhgap9*/ *Gli1* and *Lcp1* respectively. We included a predicted track in the IGV screenshot that indicates high-affinity binding regions for the TF that are represented by a tick and a black box surrounding it. In (c), we can see predicted *Elf5* peaks near *Arhgap9* and *Gli1*. ChIP-Atlas and the experimental TF ChIP-seq data substantiate the prediction near *Arhgap9*. Experimental data of *Elf5* back up the predictions near *Gli1*. We can also observe upregulated Pol2 activity in L1 in this area. In (d) we can see multiple predictions of *Elf5* bindings near *Lcp1*. ChIP-Atlas and the experimental TF ChIP-seq data corroborate the bindings of *Elf5* in this area. We also observe an upregulated Pol2 activity in time points L1 and L10 in this area.

#### Supplementary Figure 5: IGV screenshot of ELF5 target genes ARHGAP39, ARHGEF2, and IGFALS

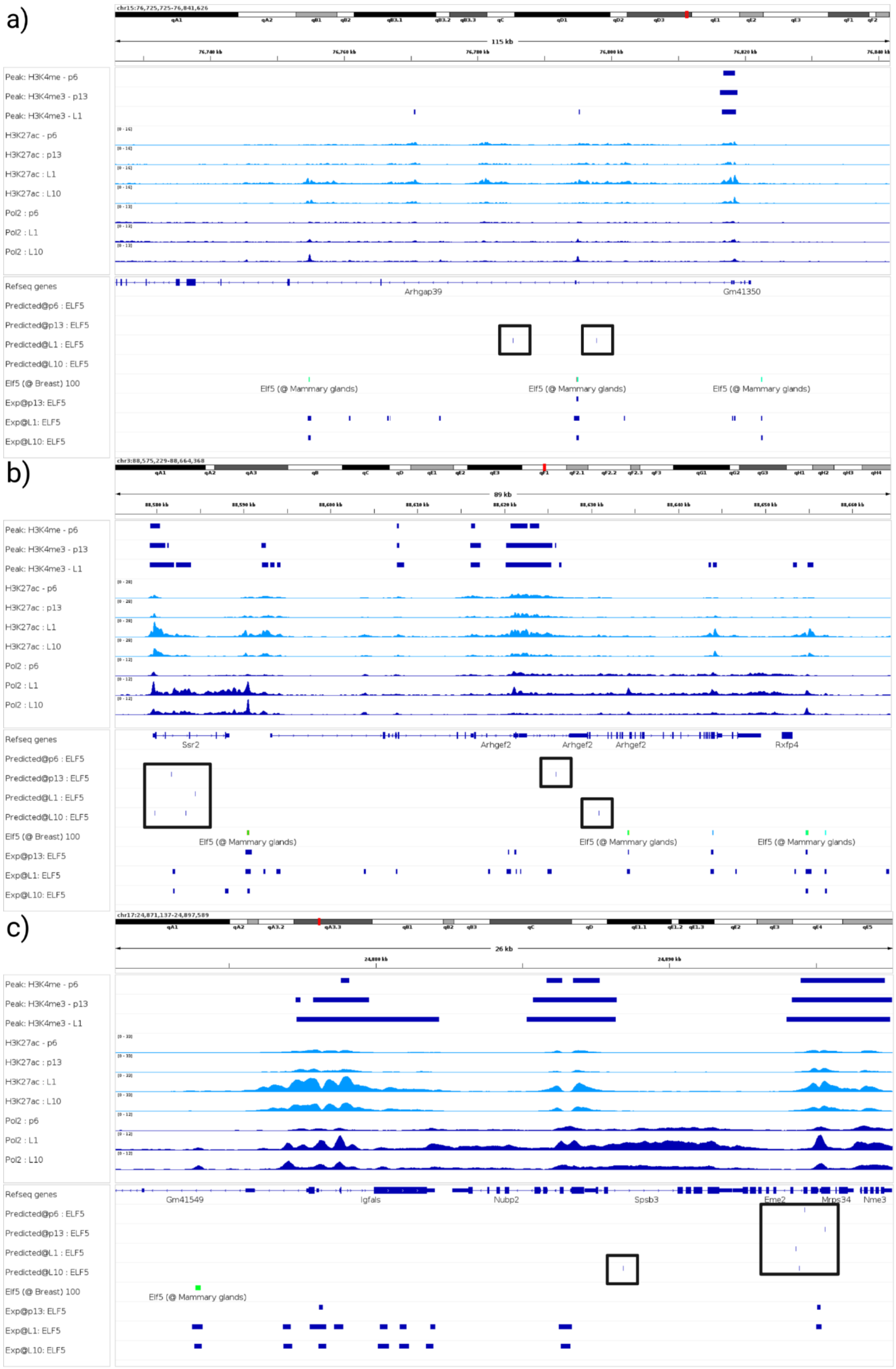

**Supplementary Figure 5:** We showcase three more examples of predicted peaks to experimental data for the transcription factor ELF5. With our predictions, we found two more genes associated with the Rho GTPase (ARHGAP39, ARHGEF2) and IGFALS that are known to play a role in mammary gland development and lactation (see Suppl. Material 2 for detailed discussion).

#### Supplementary Figure 6: Co-occurrence analysis

a) Relative Overlap

| | ARNT | CREB1 | ELF5 | ESR1 | KLF4 | MYC | NFIB | SNAI2 | STAT1 | STAT5A | STAT5B | TFAP2C |
|---|---|---|---|---|---|---|---|---|---|---|---|---|
| ARNT | - | 0.46 | 0.29 | 0.7 | 0.01 | 0.09 | 0.41 | 0.06 | 0.03 | 0.42 | 0.11 | 0.21 |
| CREB1 | 0.46 | - | 0.54 | 0.24 | - | 0.04 | 0.73 | 0.02 | 0.01 | 0.82 | 0.17 | 0.24 |
| ELF5 | 0.29 | 0.54 | - | 0.34 | 0.01 | 0.05 | 0.96 | 0.04 | 0.02 | 1.87 | 0.32 | 0.06 |
| ESR1 | 0.7 | 0.24 | 0.34 | - | 0.01 | 0.08 | 0.38 | 0.06 | 0.03 | 0.65 | 0.1 | 0.12 |
| KLF4 | 0.01 | - | 0.01 | 0.01 | - | 0.3 | 0.01 | 0.02 | 0.44 | 0.02 | 0.01 | - |
| MYC | 0.09 | 0.04 | 0.05 | 0.08 | 0.3 | - | 0.04 | 0.1 | 0.44 | 0.06 | 0.07 | 0.02 |
| NFIB | 0.41 | 0.73 | 0.96 | 0.38 | 0.01 | 0.04 | - | 0.03 | 0.01 | 1.7 | 0.24 | 0.14 |
| SNAI2 | 0.06 | 0.02 | 0.04 | 0.06 | 0.02 | 0.1 | 0.03 | - | 0.04 | 0.03 | 0.05 | 0.03 |
| STAT1 | 0.03 | 0.01 | 0.02 | 0.03 | 0.44 | 0.44 | 0.01 | 0.04 | - | 0.03 | 0.03 | - |
| STAT5A | 0.42 | 0.82 | 1.87 | 0.65 | 0.02 | 0.06 | 1.7 | 0.03 | 0.03 | - | 0.98 | 0.06 |
| STAT5B | 0.11 | 0.17 | 0.32 | 0.1 | 0.01 | 0.07 | 0.24 | 0.05 | 0.03 | 0.98 | - | 0.02 |
| TFAP2C | 0.21 | 0.24 | 0.06 | 0.12 | - | 0.02 | 0.14 | 0.03 | - | 0.06 | 0.02 | - |

b) Log-likelihood Score

| | ARNT | CREB1 | ELF5 | ESR1 | KLF4 | MYC | NFIB | SNAI2 | STAT1 | STAT5A | STAT5B | TFAP2C |
|---|---|---|---|---|---|---|---|---|---|---|---|---|
| ARNT | | 667.49 | 73.28 | 60.23 | 118.75 | 293.18 | 31.59 | 6.61 | 60.25 | 44.70 | 21.86 | 15.58 |
| CREB1 | 667.49 | | 6,914.01 | 3,856.06 | 26.57 | 330.27 | 15,396.66 | 383.39 | 11.57 | 5,902.30 | 3,199.86 | 344.21 |
| ELF5 | 73.28 | 6,914.01 | | 9,269.84 | 51.88 | 74.98 | 7,172.26 | 302.60 | 19.35 | 8,750.11 | 5,136.57 | 3,395.72 |
| ESR1 | 60.23 | 3,856.06 | 9,269.84 | | 29.61 | 35.56 | 8,492.46 | 1,210.72 | 0.84 | 8,464.57 | 1,360.74 | 6,445.27 |
| KLF4 | 118.75 | 26.57 | 51.88 | 29.61 | | 212.27 | 49.79 | 125.35 | 73.15 | 54.65 | 109.25 | 53.05 |
| MYC | 293.18 | 330.27 | 74.98 | 35.56 | 212.27 | | 147.37 | 171.35 | 156.47 | 66.42 | 158.91 | 261.56 |
| NFIB | 31.59 | 15,396.66 | 7,172.26 | 8,492.46 | 49.79 | 147.37 | | 401.70 | 12.47 | 5,237.32 | 3,015.95 | 1,192.97 |
| SNAI2 | 6.61 | 383.39 | 302.60 | 1,210.72 | 125.35 | 171.35 | 401.70 | | 92.37 | 323.40 | 15.93 | 7.65 |
| STAT1 | 60.25 | 11.57 | 19.35 | 0.84 | 73.15 | 156.47 | 12.47 | 92.37 | | 21.48 | 76.50 | 9.37 |
| STAT5A | 44.70 | 5,902.30 | 8,750.11 | 8,464.57 | 54.65 | 66.42 | 5,237.32 | 323.40 | 21.48 | | 12,299.78 | 3,822.40 |
| STAT5B | 21.86 | 3,199.86 | 5,136.57 | 1,360.74 | 109.25 | 158.91 | 3,015.95 | 15.93 | 76.50 | 12,299.78 | | 555.05 |
| TFAP2C | 15.58 | 344.21 | 3,395.72 | 6,445.27 | 53.05 | 261.56 | 1,192.97 | 7.65 | 9.37 | 3,822.40 | 555.05 | |

b) Sample size

| | ARNT | CREB1 | ELF5 | ESR1 | KLF4 | MYC | NFIB | SNAI2 | STAT1 | STAT5A | STAT5B | TFAP2C |
|---|---|---|---|---|---|---|---|---|---|---|---|---|
| ARNT | 10578 | 3791 | 1508 | 3693 | 56 | 200 | 2856 | 395 | 49 | 1500 | 622 | 2714 |
| CREB1 | 3791 | 38224 | 16531 | 7622 | 42 | 318 | 23923 | 380 | 46 | 15180 | 5216 | 11002 |
| ELF5 | 1508 | 16531 | 36223 | 3280 | 53 | 181 | 18451 | 414 | 52 | 16581 | 6076 | 2609 |
| ESR1 | 3693 | 7622 | 3280 | 77266 | 58 | 234 | 6130 | 590 | 52 | 3150 | 1126 | 7394 |
| KLF4 | 56 | 42 | 53 | 58 | 65 | 45 | 56 | 51 | 16 | 53 | 51 | 57 |
| MYC | 200 | 318 | 181 | 234 | 45 | 367 | 253 | 121 | 40 | 168 | 132 | 307 |
| NFIB | 2856 | 23923 | 18451 | 6130 | 56 | 253 | 45079 | 507 | 52 | 16263 | 5521 | 6521 |
| SNAI2 | 395 | 380 | 414 | 590 | 51 | 121 | 507 | 5897 | 52 | 348 | 361 | 1238 |
| STAT1 | 49 | 46 | 52 | 52 | 16 | 40 | 52 | 52 | 111 | 52 | 52 | 48 |
| STAT5A | 1500 | 15180 | 16581 | 3150 | 53 | 168 | 16263 | 348 | 52 | 34000 | 8265 | 1985 |
| STAT5B | 622 | 5216 | 6076 | 1126 | 51 | 132 | 5521 | 361 | 52 | 8265 | 8502 | 743 |
| TFAP2C | 2714 | 11002 | 2609 | 7394 | 57 | 307 | 6521 | 1238 | 48 | 1985 | 743 | 44526 |

**Supplementary Figure 6:** We show the co-occurrence analysis of prioritized TFs. We can see that CREB1 has a high overlap of peaks with ELF5, NFIB, STAT5A, and STAT5b which are all key players in mammary gland development and lactation.

#### Supplementary Figure 7: Co-occurrence analysis

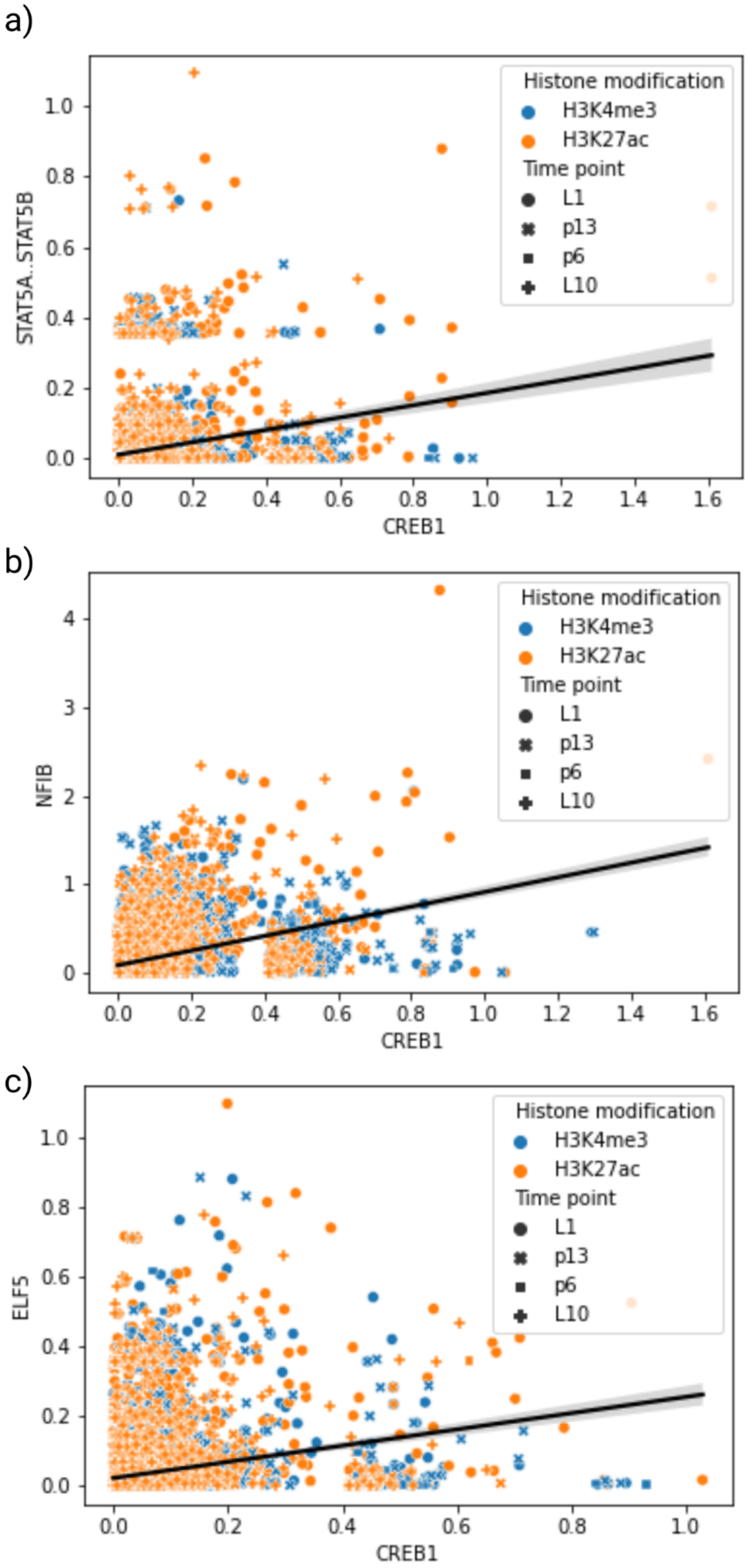

**Supplementary Figure 7:** We show the binding sites that co-occur between CREB1 and STAT5A..STAT5B, NFIB, or ELF5. We can see that there is a positive trend between the TFs. IF CREB1 has a higher binding affinity, the other TF that co-occurs on the same binding site also has a higher binding affinity on average.

**Supplementary Figure 8: Running TF-Prioritizer on open-chromatin peaks of ATAC-seq and DNase-seq instead of footprints that are corrected for protocol-specific biases**

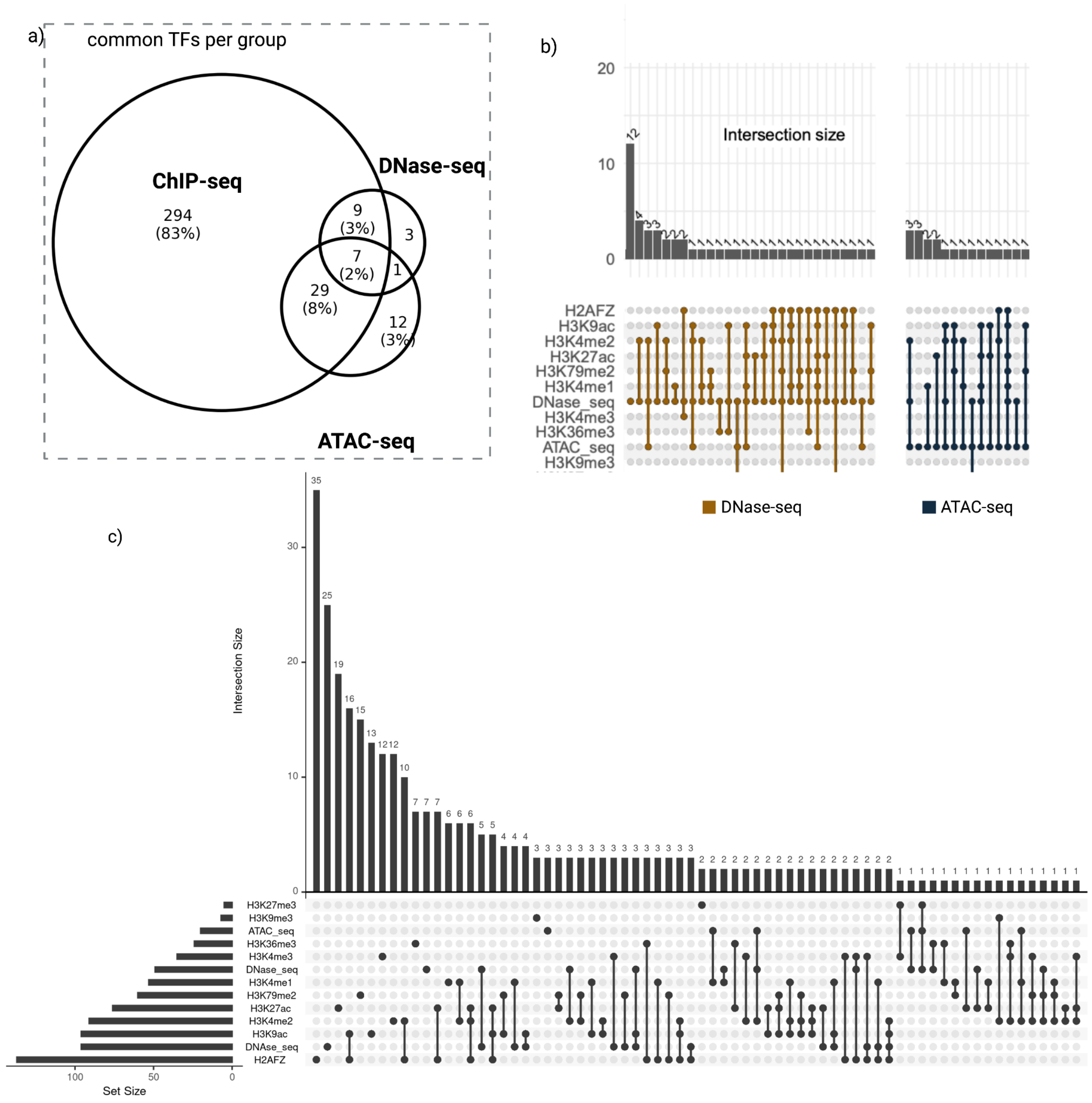

**Supplementary Figure 8:** The above plots describe the common TFs across the different methods, ATAC-seq, DNase-seq, and ChIP-seq histone modifications, without correcting for the technical biases between the protocols using HINT. a) shows the overlapping TFs between ChIP-seq, ATAC-seq, and DNase-seq independent of single histone modifications. b) displays individual intersections of TFs between all possible combinations grouped by ATAC- and DNase-seq. c) represents ungrouped intersections between groups of the first X biggest overlaps.

**Supplementary Figure 9: Comparisons of overlaps of open-chromatin peaks of ATAC-seq, DNase-seq, and ChIP-seq and overlaps of footprints of ATAC-seq and DNase-seq that were corrected for protocol-specific biases with open-chromatin peaks in ChIP-seq**

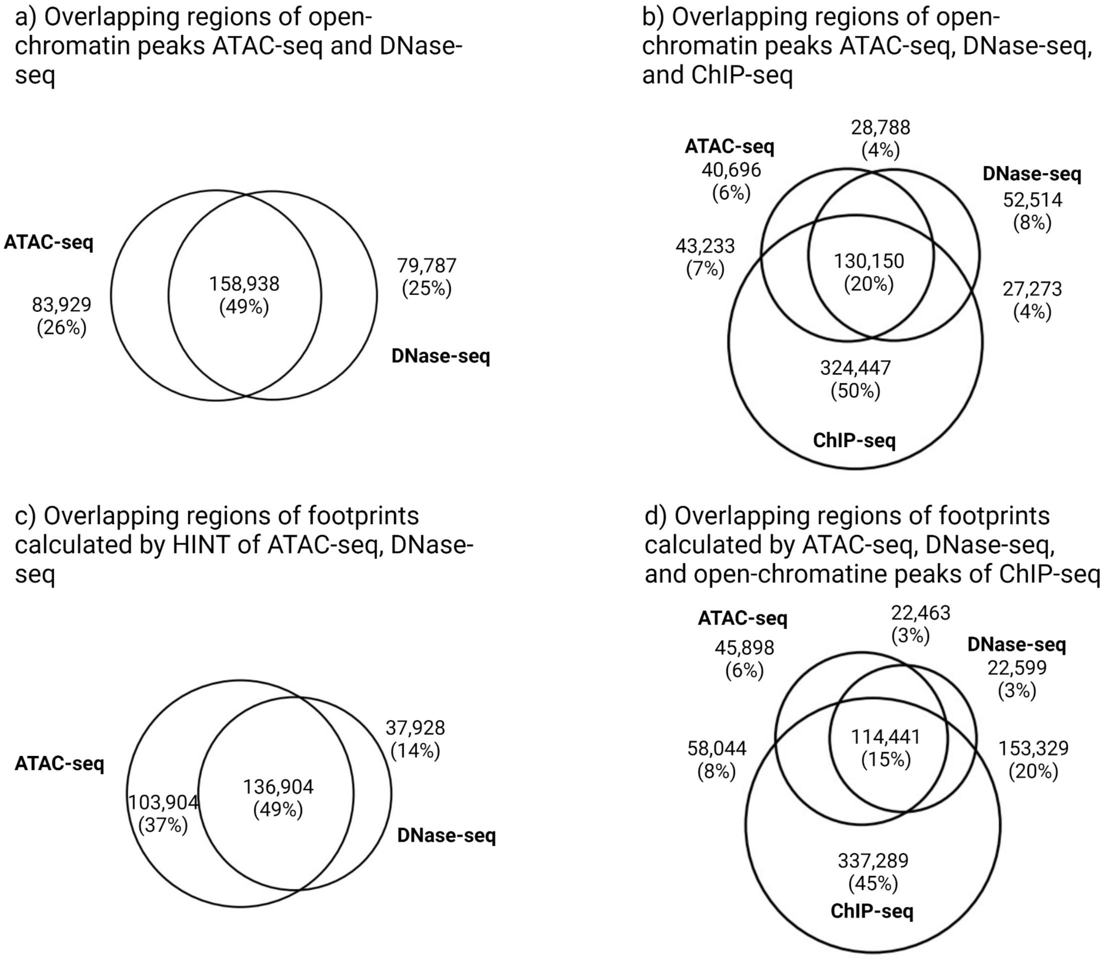

**Supplementary Figure 9:** a) Analysis of overlaps between open-chromatin peaks in ATAC-seq and DNase-seq. b) Analysis of overlaps between open-chromatin peaks in ATAC-seq, DNase-seq, and ChIP-seq. c) Analysis of protocol bias-corrected footprints between ATAC-seq and DNase-seq. d) Analysis of protocol bias-corrected footprints between ATAC-seq, DNase-seq, and open-chromatin peaks of ChIP-seq.

#### Supplementary Figure 10:

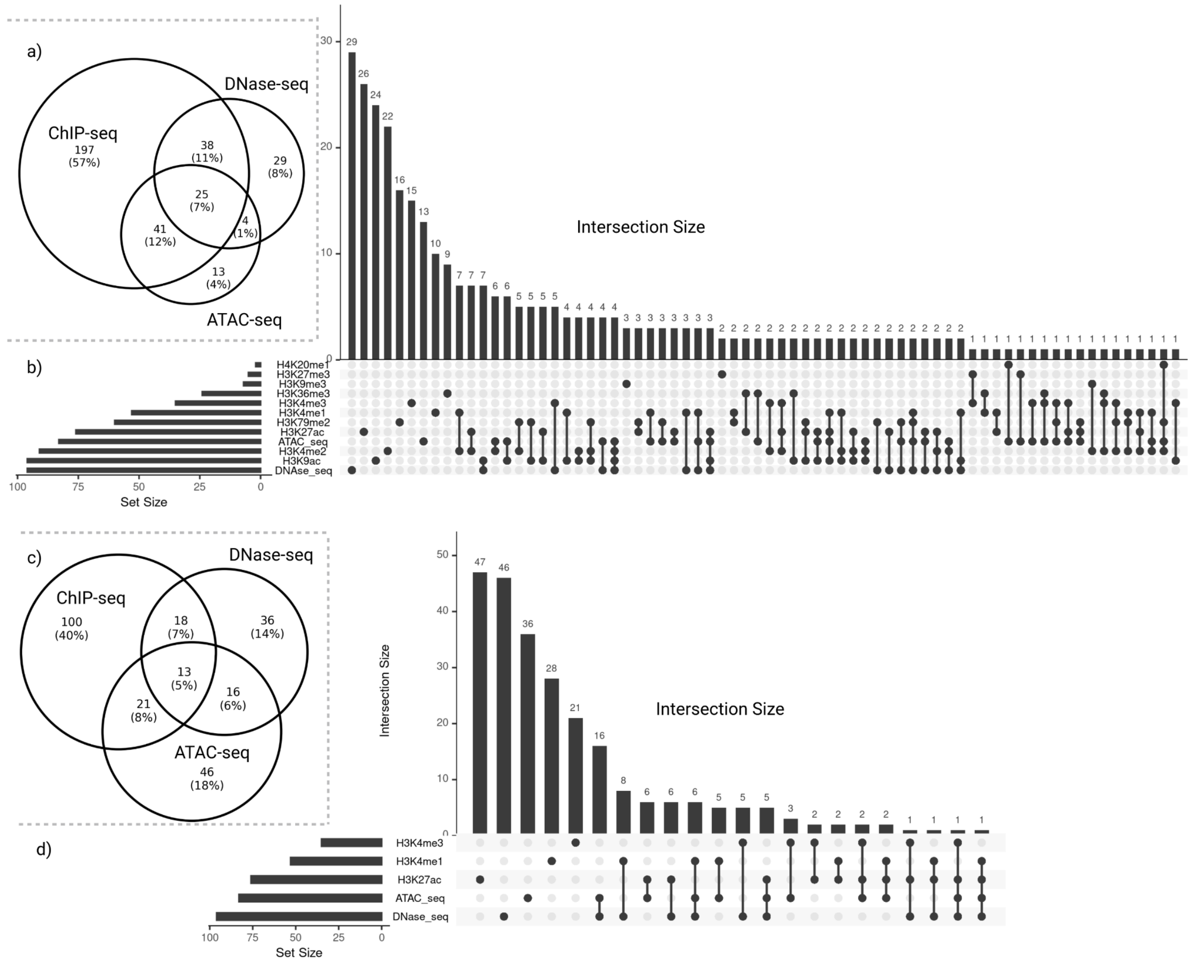

**Supplementary Figure 10:** a) and b) Due to the high number of TFs found in H2AFZ, we excluded this histone variant. We can see that the number of identified TFs dropped from a total of 339 to 301. However, it also excluded some TFs that were identified by ATAC-seq and DNase-seq. c) and d) shows how the number of identified TFs behave if one only includes the frequently used HM ChIP-seq data H3K4me3, H3K4me1, and H3K27ac in comparison to DNase-seq and ATAC-seq. We can see a drop in totally identified TFs from 339 to 152 in ChIP-seq. However, also the number of overlaps between ATAC-seq and DNase-seq drops.

**Supplementary Material 1: ENCODE file identifiers**

a) ATAC-seq BAM files
   i) K562

ENCSR483RKN_1.bam ENCSR483RKN_2.bam.bai ENCSR859USB_2.bam
ENCSR868FGK_1.bam.bai ENCSR868FGK_3.bam ENCSR956DNB_1.bam.bai
ENCSR483RKN_1.bam.bai ENCSR859USB_1.bam ENCSR859USB_2.bam.bai
ENCSR868FGK_2.bam ENCSR868FGK_3.bam.bai ENCSR956DNB_2.bam
ENCSR483RKN_2.bam ENCSR859USB_1.bam.bai ENCSR868FGK_1.bam
ENCSR868FGK_2.bam.bai ENCSR956DNB_1.bam ENCSR956DNB_2.bam.bai

   ii) MCF-7

ENCSR422SUG_1.bam ENCSR422SUG_1.bam.bai ENCSR422SUG_2.bam
ENCSR422SUG_2.bam.bai

b) ATAC-seq narrow peak bed files
   i) K562

ENCSR483RKN_1.bed ENCSR483RKN_2.bed ENCSR859USB_1.bed
ENCSR859USB_2.bed ENCSR868FGK_1.bed ENCSR868FGK_2.bed
ENCSR868FGK_3.bed ENCSR956DNB_1.bed ENCSR956DNB_2.bed

   ii) MCF-7

ENCSR422SUG_1.bed ENCSR422SUG_2.bed

c) DNase-seq BAM files
   i) K562

ENCSR000EKP_1.bam ENCSR000EKQ_1.bam ENCSR000EKS_1.bam
ENCSR000EOT_1.bam ENCSR000EOT_2.bam

   ii) MCF-7

ENCSR000EKV_1.bam ENCSR000EKW_1.bam ENCSR000EKZ_1.bam
ENCSR000EPH_1.bam ENCSR000EPH_2.bam

d) DNase-seq narrow peak bed files
   i) K562

ENCSR000EKP_1.bed ENCSR000EKQ_1.bed ENCSR000EKS_1.bed
ENCSR000EOT_1.bed ENCSR000EOT_2.bed

   ii) MCF-7

ENCSR000EKV_1.bed ENCSR000EKW_1.bed ENCSR000EKZ_1.bed
ENCSR000EPH_1.bed ENCSR000EPH_2.bed

e) HM ChIP-seq broad peak files
   i) H2AFZ
      1) K562

ENCFF000BWU_H2AFZ_REP1.mLb.clN_peaks.broadPeak
ENCFF000BWV_H2AFZ_REP1.mLb.clN_peaks.broadPeak

      2) MCF-7

ENCFF081DDG_H2AFZ_REP1.mLb.clN_peaks.broadPeak
ENCFF565MIO_H2AFZ_REP1.mLb.clN_peaks.broadPeak

ii) H3K27ac

1) K562

ENCFF000BXG_H3K27ac_REP1.mLb.clN_peaks.broadPeak
ENCFF000BXH_H3K27ac_REP1.mLb.clN_peaks.broadPeak
ENCFF143EVF_H3K27ac_REP1.mLb.clN_peaks.broadPeak
ENCFF845TAO_H3K27ac_REP1.mLb.clN_peaks.broadPeak

2) MCF-7

ENCFF000VHC_H3K27ac_REP1.mLb.clN_peaks.broadPeak
ENCFF000VHD_H3K27ac_REP1.mLb.clN_peaks.broadPeak
ENCFF303DDV_H3K27ac_REP1.mLb.clN_peaks.broadPeak
ENCFF787RSQ_H3K27ac_REP1.mLb.clN_peaks.broadPeak

iii) H3K27me3

1) K562

ENCFF000BXN_H3K27me3_REP1.mLb.clN_peaks.broadPeak
ENCFF000VDN_H3K27me3_REP1.mLb.clN_peaks.broadPeak
ENCFF699LBD_H3K27me3_REP1.mLb.clN_peaks.broadPeak
ENCFF000BXP_H3K27me3_REP1.mLb.clN_peaks.broadPeak
ENCFF000VDP_H3K27me3_REP1.mLb.clN_peaks.broadPeak
ENCFF936TRH_H3K27me3_REP1.mLb.clN_peaks.broadPeak

2) MCF-7

ENCFF000VFU_H3K27me3_REP1.mLb.clN_peaks.broadPeak
ENCFF000VFY_H3K27me3_REP1.mLb.clN_peaks.broadPeak
ENCFF134NCG_H3K27me3_REP1.mLb.clN_peaks.broadPeak
ENCFF738ERP_H3K27me3_REP1.mLb.clN_peaks.broadPeak
ENCFF000VFW_H3K27me3_REP1.mLb.clN_peaks.broadPeak
ENCFF023EPH_H3K27me3_REP1.mLb.clN_peaks.broadPeak
ENCFF250WAH_H3K27me3_REP1.mLb.clN_peaks.broadPeak

iv) H3K36me3

1) K562

ENCFF000BXO_H3K36me3_REP1.mLb.clN_peaks.broadPeak
ENCFF001FWV_H3K36me3_REP1.mLb.clN_peaks.broadPeak
ENCFF544IFT_H3K36me3_REP1.mLb.clN_peaks.broadPeak
ENCFF000BXR_H3K36me3_REP1.mLb.clN_peaks.broadPeak
ENCFF001FWW_H3K36me3_REP1.mLb.clN_peaks.broadPeak
ENCFF627AZR_H3K36me3_REP1.mLb.clN_peaks.broadPeak

2) MCF-7

ENCFF000VGE_H3K36me3_REP1.mLb.clN_peaks.broadPeak
ENCFF000VGI_H3K36me3_REP1.mLb.clN_peaks.broadPeak
ENCFF082PFU_H3K36me3_REP1.mLb.clN_peaks.broadPeak
ENCFF389PKW_H3K36me3_REP1.mLb.clN_peaks.broadPeak
ENCFF000VGG_H3K36me3_REP1.mLb.clN_peaks.broadPeak
ENCFF055OLK_H3K36me3_REP1.mLb.clN_peaks.broadPeak

ENCFF147WQG_H3K36me3_REP1.mLb.clN_peaks.broadPeak
ENCFF819ITL_H3K36me3_REP1.mLb.clN_peaks.broadPeak

v) H3K4me1

1) K562

ENCFF000BXX_H3K4me1_REP1.mLb.clN_peaks.broadPeak
ENCFF000VDU_H3K4me1_REP1.mLb.clN_peaks.broadPeak
ENCFF399XNM_H3K4me1_REP1.mLb.clN_peaks.broadPeak
ENCFF000BYG_H3K4me1_REP1.mLb.clN_peaks.broadPeak
ENCFF000VDV_H3K4me1_REP1.mLb.clN_peaks.broadPeak
ENCFF646KOA_H3K4me1_REP1.mLb.clN_peaks.broadPeak

2) MCF-7

ENCFF083FLF_H3K4me1_REP1.mLb.clN_peaks.broadPeak
ENCFF415RHX_H3K4me1_REP1.mLb.clN_peaks.broadPeak
ENCFF522RPS_H3K4me1_REP1.mLb.clN_peaks.broadPeak
ENCFF197FAY_H3K4me1_REP1.mLb.clN_peaks.broadPeak
ENCFF461WBE_H3K4me1_REP1.mLb.clN_peaks.broadPeak

vi) H3K4me2

1) K562

ENCFF000BYA_H3K4me2_REP1.mLb.clN_peaks.broadPeak
ENCFF000BYF_H3K4me2_REP1.mLb.clN_peaks.broadPeak

2) MCF-7

ENCFF002ACV_H3K4me2_REP1.mLb.clN_peaks.broadPeak
ENCFF002AUN_H3K4me2_REP1.mLb.clN_peaks.broadPeak

vii) H3K4me3

1) K562

ENCFF000BYI_H3K4me3_REP1.mLb.clN_peaks.broadPeak
ENCFF000VDX_H3K4me3_REP1.mLb.clN_peaks.broadPeak
ENCFF001FXG_H3K4me3_REP1.mLb.clN_peaks.broadPeak
ENCFF000BYJ_H3K4me3_REP1.mLb.clN_peaks.broadPeak
ENCFF000VDY_H3K4me3_REP1.mLb.clN_peaks.broadPeak
ENCFF001FXH_H3K4me3_REP1.mLb.clN_peaks.broadPeak

2) MCF-7

ENCFF001FYE_H3K4me3_REP1.mLb.clN_peaks.broadPeak
ENCFF020LCP_H3K4me3_REP1.mLb.clN_peaks.broadPeak
ENCFF298UCU_H3K4me3_REP1.mLb.clN_peaks.broadPeak
ENCFF792HDC_H3K4me3_REP1.mLb.clN_peaks.broadPeak
ENCFF001FYK_H3K4me3_REP1.mLb.clN_peaks.broadPeak
ENCFF211APE_H3K4me3_REP1.mLb.clN_peaks.broadPeak
ENCFF768PAZ_H3K4me3_REP1.mLb.clN_peaks.broadPeak
ENCFF923ABZ_H3K4me3_REP1.mLb.clN_peaks.broadPeak

viii) H3K79me2

1) K562

ENCFF000BYO_H3K79me2_REP1.mLb.clN_peaks.broadPeak
ENCFF000BYS_H3K79me2_REP1.mLb.clN_peaks.broadPeak

ENCFF296QNV_H3K79me2_REP1.mLb.clN_peaks.broadPeak
ENCFF946WKP_H3K79me2_REP1.mLb.clN_peaks.broadPeak

2) MCF-7

ENCFF004XWK_H3K79me2_REP1.mLb.clN_peaks.broadPeak
ENCFF022HZP_H3K79me2_REP1.mLb.clN_peaks.broadPeak
ENCFF161ZED_H3K79me2_REP1.mLb.clN_peaks.broadPeak
ENCFF402WSG_H3K79me2_REP1.mLb.clN_peaks.broadPeak

ix) H3K9ac

1) K562

ENCFF000BYW_H3K9ac_REP1.mLb.clN_peaks.broadPeak
ENCFF000VEE_H3K9ac_REP1.mLb.clN_peaks.broadPeak
ENCFF000VEG_H3K9ac_REP1.mLb.clN_peaks.broadPeak
ENCFF001QWV_H3K9ac_REP1.mLb.clN_peaks.broadPeak

2) MCF-7

ENCFF440EIF_H3K9ac_REP1.mLb.clN_peaks.broadPeak
ENCFF564IMY_H3K9ac_REP1.mLb.clN_peaks.broadPeak
ENCFF815TBB_H3K9ac_REP1.mLb.clN_peaks.broadPeak
ENCFF921PCJ_H3K9ac_REP1.mLb.clN_peaks.broadPeak

x) H3K9me3

1) K562

ENCFF001QWW_H3K9me3_REP1.mLb.clN_peaks.broadPeak
ENCFF001QWX_H3K9me3_REP1.mLb.clN_peaks.broadPeak

2) MCF-7

ENCFF000VFE_H3K9me3_REP1.mLb.clN_peaks.broadPeak
ENCFF000VFJ_H3K9me3_REP1.mLb.clN_peaks.broadPeak
ENCFF600JOS_H3K9me3_REP1.mLb.clN_peaks.broadPeak
ENCFF000VFG_H3K9me3_REP1.mLb.clN_peaks.broadPeak
ENCFF517BLK_H3K9me3_REP1.mLb.clN_peaks.broadPeak
ENCFF656BIN_H3K9me3_REP1.mLb.clN_peaks.broadPeak

xi) H4K20me1

1) K562

ENCFF000BZJ_H4K20me1_REP1.mLb.clN_peaks.broadPeak
ENCFF000BZN_H4K20me1_REP1.mLb.clN_peaks.broadPeak

2) MCF-7

ENCFF465RWE_H4K20me1_REP1.mLb.clN_peaks.broadPeak
ENCFF475AEQ_H4K20me1_REP1.mLb.clN_peaks.broadPeak
ENCFF476VWW_H4K20me1_REP1.mLb.clN_peaks.broadPeak
ENCFF528WDJ_H4K20me1_REP1.mLb.clN_peaks.broadPeak

**Supplementary Material 2: Confusion matrices and the calculation of sensitivity, specificity, precision, accuracy, and F1-score**

a) Confusion Matrix

| | Experimental | |
|---|---|---|
| Predicted | True | False |
| True | TP | FP |
| False | FN | TN |

TP = True Positives
FP = False Positives
TN = True Negatives
FN = False Negatives

b) Metrics

Sensitivity = $\frac{TP}{TP+FN}$

Specificity = $\frac{TN}{TN+FP}$

Precision = $\frac{TP}{TP+FP}$

Accuracy = $\frac{TP+TN}{TP+FP+TN+FN}$

F1-Score = $\frac{2 \text{ x Sensitivity x Precision}}{\text{Sensitivity + Precision}}$

#### Supplementary Material 3: Biological findings

In our application case (https://exbio.wzw.tum.de/tfprio/mouse/#/), TF-Prioritizer reports several TFs known to be involved in mammary gland development and/or lactation, including Signal Transducer and Activator of Transcription (STAT5 - consisting of STAT5A and STAT5B) [16–18], E74 Like ETS Transcription Factor 5 (ELF5) [19,20], Estrogen Receptor 1 (ESR1) [21], and Nuclear Factor I B (NFIB) [18]. TF-Prioritizer also identifies TFs that are known to be important in pregnancy, e.g., ETS Proto-Oncogene 2 (ETS2) [22]. Furthermore, we prioritize a few candidate TFs that are not yet widely known to be involved in either of the processes (e.g., CAMP Responsive Element Binding Protein 1 (CREB1), Aryl Hydrocarbon Receptor Nuclear Translocator (ARNT)) showing the potential of TF-Prioritizer to generate new hypotheses, e.g., overall, we found that 94 out of 104 prioritized TFs controlled at least one Rho family GTPase-associated target gene. Rho family GTPases play an important role in epithelial morphogenesis during mammary gland development [23,24]. Furthermore, we predict 58 of 104 prioritized TFs to control genes of the Casein (*Csn*) family proteins that are known to be milk proteins [25].

In the following, we intensively evaluate and discuss the TFs STAT5 and ELF5 and their predicted target genes, as those TFs are widely accepted to be important in mammary gland development and lactation in mice. We investigate for each TF its expression change (DESeq2 normalized gene counts) over the time points (pregnancy day 6 (p6), day 13 (p13), and lactation day 1 (L1), day 10 (L10)) as well as the top 30 predicted target genes for selected histone modifications and time points. We evaluate the sensitivity, specificity, precision, and accuracy of the predicted peaks using experimentally validated data to review the literature about the TF's role in pregnancy, mammary gland development, or lactation. We pick a few of the predicted target genes for closer evaluation and assess differential expression between the two stages (pregnancy and lactation). We determine if we predict the high binding energy of the TF in close proximity to the target gene and evaluate the

predicted peaks - the position of the highest affinity value of the TF in the open chromatin region predicted by TRAP - with experimental evidence using external data from ChIP-Atlas and experimental data from pregnant and lactating mice. Also, we evaluate the expression change of the target gene on a Pol2 ChIp-seq signal. Lastly, we provide literature and interpretation of the target gene's role in pregnancy, mammary gland development, or lactation.
After the evaluation of STAT5 and ELF5, we provide further details about the predictions of the TFs ESR1, NFIB, CREB1, and ARNT with respect to the Rho family GTPase-associated target genes and the Casein protein family. For further data exploration, we refer to our web application for all prioritized TFs (see the Availability section).

#### STAT5

*Stat5* mRNA levels are highly upregulated during the last days of pregnancy and at the beginning of lactation from p6 (1,076), p13 (2,810) to L1 (3,355), and L10 (1,203). In Figure 3. a) (H3K4me3) and 3b (H3K27ac), we can see a clear expression separation between p13 and L1 of predicted target genes of STAT5. Suppl. Fig. 2. a) shows that TF-Prioritizer is able to reach a sensitivity of 57.89%, a specificity of 66.39%, a precision of 78.15%, an accuracy of 60.65%, and an F1 score of 66.51% for STAT5A..STAT5B. These high percentages of statistical measures give us confidence in our predictions. STAT5 is known in the literature to significantly regulate mammary gland morphology [26].
Suppressor Of Cytokine Signaling 2 (*Socs2*) (Figure 3. a) mRNA levels show higher expression in pregnancy compared to lactation. *Socs2* shows both experimental and predicted peaks (Figure 3. c). We can also observe a change in the Pol2 signal between pregnancy and lactation, reflecting the increased transcriptional activity. SOCS2 has distinct physiological functions in the developing mammary gland [27], and STAT5 may act as a regulator for SOCS2 during pregnancy and lactation [28].
mRNA levels of the members of the Casein (*Csn*) family are strongly upregulated during lactation (Figure 3. b). This includes the genes of the milk protein Casein Beta (*Csn2*) [29] and the Casein Alpha S2 Like proteins (*Csn1s2a* and *Csn1s2b*) [30,31]. STAT5 ChIP-Atlas data supports the binding of predicted STAT5 peaks in the genomic area surrounding these genes. We can also observe a precise change in TF binding activity during p6 (few peaks), p13 (more peaks), and L1 (crowded peaks) using complementary STAT5 experimental TF ChIP-seq data from lactating mice (Figure 3. d). In Pol2 signaling (Figure 3. d), we observe a significant change between p6 and L1. mRNA levels of Discoidin Domain Receptor Tyrosine Kinase 1 (*Ddr1*) appears to be upregulated by STAT5 (Figure 3. a-b). In Suppl. Fig. 3, we observe that TRAP predicted high binding energy for STAT5 to the regulatory region of *Ddr1* in L1. Experimental data of ChIP-Atlas and STAT5 ChIP-seq data also indicate the binding of STAT5 in the *Ddr1* region, confirming our predicted peak. We can also observe a significant increase in the Pol2 signal during the time between p6 and L1 in the predicted region. STAT5 is known to be linked to *Ddr1,* which is essential in mammary gland development [32,33], as DDR1 signaling is essential to sustain STAT5 function during lactogenesis [34].

#### ELF5

Past studies have shown that ELF5 is important for mammary gland development [19,20]. Indeed, *Elf5* mRNA levels show increasing expression at the beginning of the pregnancy p6 (1,355), p13 (6,970) to lactation L1 (12,729), L10 (6,133) (Figure 4. a-b). The heatmap of the top 30 predicted target genes in Figure 4. a) (H3K4me3 - p6 versus L1) and Figure 4. b) (Pol2 - p6 versus L10) clearly separates predicted target genes between pregnancy and lactation. TF-Prioritizer can predict peaks for ELF5 with a sensitivity of 77.57%, a specificity of 80.59%, a precision of 81.59%, an accuracy of 79.00%, and an F1 score of 79.53%, which indicates that these peaks are correctly predicted.
Figure 4. a) shows that ELF5 likely leads to the downregulation of GLI Family Zinc Finger 1 (*Gli1*) mRNA levels during lactation. In Figure 4. c), we can witness predicted and experimentally validated peaks near *Gli1*. We can observe that the Pol2 signal is increasing over time to lactation. In close proximity to *Gli1*, we can observe Rho GTPase Activating

Protein 9 (*Arhgap9*), whose mRNA levels are predicted to be upregulated by ELF5. We believe that ELF5 is acting as a suppressor for *Gli1* mRNA levels as Fiaschi et al. showed that *Gli1*-expressing females were unable to lactate, and milk protein gene expression was essentially absent [35].

Figure 4. a) further shows that Rho GTPase Activating Protein 9 (*Arhgap9*) mRNA levels, once translated ARGAP9 becomes one of several essential proteins in Rho GTPases [23,24], are upregulated in lactation compared to pregnancy. We predict high binding energy in close proximity and inside *Arhgap9*. In Figure 4. c), we observe experimentally validated peaks which are corroborated by results from ChIP-Atlas. We also notice a significant change in the Pol2 signal between pregnancy and lactation in this area.

In Figure 4. c), we observe that *Arhgap9* and *Gli1* are close neighbors in the human genome. We hypothesize that ELF5 is suppressing mRNA levels of *Gli1* to enable lactation [35] and is upregulating mRNA levels of *Arhgap9* for Rho GTPase activity during lactation at the same time.

In Figure 4. b), Rho GTPase Activating Protein 39 (*Arhgap39*) mRNA levels are upregulated during pregnancy. ARHGAP9 is another Rho GTPase activating protein and could therefore be essential for mammary gland development. Suppl. Fig. 4 shows predicted peaks close to *Arhgap39*. We also observe experimentally validated peaks which are corroborated by data from ChIP-Atlas. In addition, we notice an increase in the Pol2 signals in this area during lactation compared to pregnancy.

Rho/Rac Guanine Nucleotide Exchange Factor 2 (*Arhgef2*) mRNA levels, which are essential for Rho GTPase activity, are also upregulated during pregnancy (Figure 4. b). In Suppl. Fig. 4. a-b), predicted and experimentally validated peaks occur near *Arhgef2*. We also detect a change in the Pol2 signal during lactation.

Lymphocyte Cytosolic Protein 1 (*Lcp1*) mRNA levels, which were reported essential for lactation [36], are upregulated in lactation compared to pregnancy (Figure 4), with several predicted peaks in close proximity to *Lcp1*. We also find ELF5 experimentally validated peaks in the ChIP-Atlas data and ELF5 TF ChIP-seq data at the same position as our predicted peaks. We can also observe a stronger Pol2 signal during lactation compared to pregnancy.

Insulin-Like Growth Factor Binding Protein Acid Labile Subunit (*Igfals*) mRNA levels are upregulated in lactation [37] (Figure 4. a) with high binding affinity near *Igfals* in Suppl. Fig. 4. c). We also find experimentally validated peaks near *Igfals,* which are corroborated by data from the ChIP-Atlas. We also observe a stronger Pol2 signal during lactation than in pregnancy. *Igfals* is a regulator of the tumor suppressor protein *p53* [38] activity, and *Igfals* may thus be a protective factor preventing breast cancer in mammary gland development.

#### ESR1

*Esr1* mRNA levels are upregulated the most at time points p13 (p6 (639), p13 (2,981), L1 (742), and L10 (806)). ESR1 peaks (Suppl. Fig. 2. c) can be predicted with a sensitivity of 58.79%, a specificity of 78.93%, a precision of 90.18%, an accuracy of 63.48%, and an F1 score of 71.18%. We predict that ESR1 controls the mRNA expression of at least three Rho family GTPase-associated proteins: *Arhgap39*, Rho/Rac Guanine Nucleotide Exchange Factor 18 (*Arhgef18*), and Rho Guanine Nucleotide Exchange Factor 40 (*Arhgef40*). We also observed Casein Kinase 1 Epsilon (*Csnk1e*), a member of the Casein family, to be controlled by ESR1. From these results, we can hypothesize that ESR1 could thus play a role during mammary gland development [21]. In the literature, we found that Mueller et al. concluded that complete mammary gland development depends on the estrogen receptor, among other TFs [39]. H.L.M. Tucker et al. showed that repressing *Esr1* mRNA expression has a significant impact on mammary gland development [40].

#### NFIB

*Nfib* mRNA level expression is strongly increasing during pregnancy p6 (5,320) with the highest expression at the end of the pregnancy p13 (20,517) and decreasing during lactation L1 (8,639) to L10 (2,958). We can predict the correct peaks of NFIB with a sensitivity of 77.06%, a specificity of 86.63%, a precision of 88.64%, an accuracy of 81.13%, and an F1 score of 82.45%. We predict that NFIB regulates mRNA expression levels of Casein Kinase 2 Beta (*Csnk2b*), a member of the Casein family. According to GeneCards [41], CSNK2B is a regulatory subunit of casein kinase II/CK2. Among its related pathways in the regulation of Tumor Protein P53 (TP53) [42,43]. mRNA levels of *Csnk2b* are upregulated in lactation and could therefore play a role in tumor prevention. We predict that NFIB controls mRNA levels of Rho/Rac Guanine Nucleotide Exchange Factor 2 (*Arhgef2*), Rho Guanine Nucleotide Exchange Factor 39 (*Arhgef39*), and Cdc42 Guanine Nucleotide Exchange Factor 9 (*Arhgef9*) which are associated with Rho GTPase activity. According to our predictions, NFIB also has a partial influence on the mRNA expression of *Ddr1* (see STAT5). With respect to this data, we are in line with the currently accepted knowledge that NFIB is important for mammary gland development [18].

#### Rho GTPase's role in pregnancy, mammary gland development, and lactation

We propose that our predicted TFs regulate several Rho GTPase-associated genes as their expression changes during pregnancy and lactation. For example, we observe an upregulation of *Arhgap9* that is essential for Rho GTPase activity during lactation in comparison to pregnancy (Figure 4. a and Figure 4. c). On the other hand, *Arhgef2* is upregulated during pregnancy and downregulated during lactation. *Arhgef2* is responsible for the activity of the Rho GTPase by exchanging GDP for GTP [44]. Our data suggest mechanisms of pregnancy, mammary gland development, and lactation are dependent on Rho GTPase and its regulation by multiple TFs. Experimental validation could help to further understand those complex processes.

#### CREB1

The co-occurrence analysis of TF-Prioritizer shows that CREB1 has a high number of similar peaks with TF known to be involved in mammary gland development and lactation, namely ELF5 (22% overlap), NFIB (29% overlap), and STAT5A (21% overlap) (see Suppl. Fig. 6). *Creb1* mRNA levels are upregulated during late pregnancy from p6 (779), p13 (3,361), and early lactation L1 (1,311) to L10 (400). We hence decided to have a closer look at CREB1 and its target genes. For CREB1, TF-Prioritizer reaches a sensitivity of 82.06%, a specificity of 91.35%, a precision of 92.45%, an accuracy of 86.12%, and an F1 score of 86.94% (Suppl. Fig. 2. e). We also predict CREB1 to regulate the mRNA levels of a member of the Rho GTPase family - Rho/Rac Guanine Nucleotide Exchange Factor 18 (*Arhgef18*) and the mRNA levels of a member of the Casein protein family *Csnk1e*. To the best of our knowledge, CREB1 has not yet been widely recognized to play a role in lactation or mammary gland development. However, Yao et al. [45,46] suggest CREB1 is involved in the lactation process and regulates milk fatty acid composition in the mammary gland in goats. We recommend experimentally validating the importance of CREB1 in lactation in the mammary gland in mice.

#### ARNT

We observe a similar mRNA levels trend of *Arnt* in comparison to, e.g., *Nfib*, over time, *Arnt* mRNA levels are getting more expressed during pregnancy (p6 516, p13 2210) and are getting less expressed during lactation (L1 919, L10 283) which could mean that the TF ARNT is more involved in mammary gland development but less involved in lactation. ARNT and many other TFs can regulate gene expression using co-factors [47]. We prioritized such co-regulation with either Hypoxia Inducible Factor 1 Subunit Alpha (HIF1A) [48] or Aryl Hydrocarbon Receptor (AHR) [49]. We prioritized ARNT..HIF1A (mRNA expression of *Hif1a*: p6 (639), p13 (3,654), L1 (2,357), L10 (856)), AHR..ARNT (mRNA expression of *Ahr*: p6 (492), p13 (1,892), L1 (459), L10 (91)) and ARNT individually. We can also see that the TF partners in the complex also follow the same gene expression pattern of upregulation until the end of pregnancy and then slow downregulation during lactation. In Suppl. Fig. 2. f-g) we can see that ARNT (sensitivity: 81.76%, specificity: 60.69%, precision: 41.18%, accuracy: 66.00%, F1 score: 54.77%) has a drop if it comes to co-factoring with HIF1A (sensitivity: 57.07, specificity: 54.10%, precision: 41.89%, accuracy: 55.19%, F1 score: 48.32%) and AHR (sensitivity: 63.14%, specificity: 56.30%, precision: 40.94%, accuracy: 58.52%, F1 score: 49.68%). We want to point out that we could not retrieve experimental data for HIF1A and AHR*,* which could explain the drop in the statistical metrics. We predict that the ARNT..HIF1A complex controls the mRNA levels of Rho Guanine Nucleotide Exchange Factor 1 (*Arhgef1*), Rho GTPase Activating Protein 12 (*Arhgap12*) of the Rho GTPase family, and Casein Kinase 2 Alpha 2 (*Csnk2a2*) of the Casein protein family. In the AHR..ARNT complex, we predict mRNA levels of *Arhgef39*, *Arhgef2*, and *Arhgef40* of the Rho GTPase family to be controlled. We predict Arnt to control mRNA levels of *Csnk2a2,* a member of the Casein protein family. This could mean that ARNT could be important for the lactation process. These predictions need to be experimentally validated.

**Supplementary Table 1: Feature comparison between TEPIC2 + DYNAMITE and TF-Prioritizer**

| Feature | TF-Prioritizer | TEPIC2 + DYNAMITE |
|---|---|---|
| Includes all features of TEPIC2 + DYNAMITE | ✔ | ✔ |
| Allows automatic preprocessing of ATAC-seq and DNase-seq data using HINT | ✔ | ✘ |
| Sample combination option to reduce runtime | ✔ | ✘ |
| Filter for blacklisted genomic regions | ✔ | ✘ |
| TPM filter for TFs | ✔ | ✘ |
| TPM filter for target genes | ✔ | ✘ |
| Additional filtering layer for prioritized TFs | ✔ Background distribution model | ✘ |
| Searches automatically for outside experimental data | ✔ ChIP-Atlas | ✘ |
| Creates genome browser screenshot on loci of interest using experimental and predicted data | ✔ IGV browser | ✘ |
| Adds known Enhancers to IGV screenshots | ✔ | ✘ |
| Provide insight into predictions (e.g., show peaks in IGV) | ✔ | ✘ |
| Creates heatmaps for predicted target genes that have the highest binding affinity for each TF | ✔ | ✘ |
| Creates a feature-rich interactive web application | ✔ | ✘ |

**Supplementary Table 2: Feature comparison between TF-Prioritizer and diffTF**

| Feature | TF-Prioritizer | diffTF |
|---|---|---|
| Filter for chromosomes (e.g., sex, mitochondrial chromosomes) | ✗ | ✓ mandatory, cannot be skipped |
| Classifies TFs into activators and repressors | ✗ | ✓ |
| Sample combination option to reduce runtime | ✓ | ✗ |
| Filter for blacklisted genomic regions | ✓ | ✓ |
| TPM filter for TFs | ✓ | ✗ |
| TPM filter for target genes | ✓ | ✗ |
| Biophysical Model for TF binding sites | ✓ TRAP | ✗ Use of all binding sites |
| Probability scores for TF target gene links | ✓ TEPIC | ✗ |
| Prioritize TFs based on binding site energies,TF expression and target genes expression | ✓ Linear Regression (DYNAMITE) | ✗ |
| Additional filtering layer for prioritized TFs | ✓ Background distribution model | ✓ Background distribution model |
| Searches automatically for outside experimental data | ✓ ChIP-Atlas | ✗ |
| Creates genome browser screenshot on loci of interest using experimental and predicted data | ✓ IGV browser | ✗ |
| Adds known Enhancers to IGV screenshots | ✓ | ✗ |
| Provide insight into predictions (e.g., show peaks in IGV) | ✓ | ✗ |
| Creates heatmaps for predicted target genes that have the highest binding affinity for each TF | ✓ | ✗ |
| Creates a feature-rich interactive web application | ✓ | ✗ |

#### Supplementary Table 3: Technical comparison between TF-Prioritizer and diffTF

| Technical Features | TF-Prioritizer | diffTF |
|---|---|---|
| Provides a docker or docker-like version of the pipeline to prevent technical dependency issues | ✔ dockerized | — singularity, with dependency problems |
| Pipeline can resume if unexpected shutdown happened (e.g., server crash, process was forced to quit) | ✔ | ✘ |
| Simulation of the whole process to check for validity of parameters and files | ✔ | ✔ |
| Multi-Threading available | ✔ | ✔ |
| pipeline checks available RAM and uses less threads if not enough RAM is available | ✔ | ✘ |

**Supplementary Table 4: Comparison of prioritized transcription factors between TF-Prioritizer and diffTF**

| TF | diffTF | TF-Prioritizer |
|---|---|---|
| AHR | (x) | (x) |
| AIRE | (x) | |
| ANDR | (x) | |
| AP2B | (x) | |
| AP2C | (x) | |
| ARI3A.D | (x) | |
| ARI3A.S | (x) | |
| ARI5B | (x) | |
| ARID3A | | (x) |
| ARID5A | | (x) |
| **ARNT** | **(x)** | **(x)** |
| ARNT2 | (x) | |
| ATF1 | (x) | |
| ATF2 | (x) | |
| ATF3 | (x) | |
| BACH1 | (x) | |
| BARX2 | (x) | |
| BATF | (x) | |
| BCL6 | (x) | |
| BHE40 | (x) | |
| BHLHE40 | | (x) |
| BMAL1 | (x) | |
| BRCA1 | (x) | |
| CBFB | | (x) |
| CDC5L | (x) | |
| CEBPA | (x) | (x) |
| CEBPB | (x) | |

| | | |
|---|---|---|
| CEBPD | (x) | (x) |
| CEBPG | (x) | |
| CEBPZ | (x) | |
| CLOCK | | (x) |
| COE1 | (x) | |
| COT1.B | (x) | |
| COT1.S | (x) | |
| COT2.B | (x) | |
| COT2.S | (x) | |
| **CREB1** | **(x)** | **(x)** |
| CREM | (x) | |
| CTCF | (x) | |
| CUX1 | (x) | |
| CXXC1 | (x) | |
| DBP | (x) | (x) |
| DLX3 | (x) | |
| E2F1 | (x) | |
| E2F2 | (x) | (x) |
| E2F3 | (x) | |
| E2F4 | | (x) |
| E2F5 | (x) | |
| E2F6 | (x) | (x) |
| E2F7 | (x) | (x) |
| E4F1 | (x) | |
| EGR1 | (x) | (x) |
| EGR2 | (x) | (x) |
| EGR3 | (x) | |
| ELF1 | (x) | |
| ELF3 | (x) | (x) |
| **ELF5** | **(x)** | **(x)** |

| | | |
|---|---|---|
| ELK1 | (x) | |
| ELK3 | (x) | |
| ELK4 | (x) | (x) |
| ENOA | (x) | |
| EPAS1 | (x) | (x) |
| ERG | (x) | |
| ERR2 | (x) | |
| **ESR1** | **(x)** | **(x)** |
| **ETS1** | **(x)** | **(x)** |
| ETS2 | (x) | (x) |
| ETV4 | (x) | (x) |
| ETV5 | (x) | |
| ETV6 | | (x) |
| EVI1 | (x) | |
| FLI1 | (x) | |
| FOS | (x) | |
| FOSB | (x) | |
| FOSL2 | (x) | |
| FOXA1 | (x) | |
| FOXA3 | (x) | |
| FOXC1 | (x) | |
| FOXI1 | (x) | |
| FOXJ2 | (x) | |
| FOXJ3.A | (x) | |
| FOXJ3.S | (x) | |
| FOXM1 | (x) | |
| FOXO1 | (x) | |
| FOXO3 | (x) | (x) |
| FOXO4 | (x) | (x) |
| FOXP2 | (x) | |

| | | |
|---|---|---|
| FOXP3 | (x) | |
| FOXQ1 | (x) | |
| FUBP1 | (x) | |
| GABP1 | (x) | |
| GABPA | (x) | (x) |
| GATA2 | (x) | (x) |
| GATA3 | (x) | |
| GATA6 | (x) | |
| GCR.C | (x) | |
| GCR.S | (x) | |
| GFI1 | (x) | |
| GLI1 | (x) | |
| GLI2 | (x) | |
| GLI3 | (x) | |
| GLIS3 | (x) | |
| GMEB1 | | (x) |
| HAND1 | | (x) |
| HBP1 | (x) | |
| HES1 | (x) | (x) |
| HEY2 | (x) | |
| HIC1 | (x) | (x) |
| HIF1A | (x) | (x) |
| HINFP | (x) | |
| HLF | (x) | (x) |
| HLTF | (x) | |
| HMGA1 | (x) | |
| HNF1A | (x) | |
| HOXA5 | | (x) |
| HOXB4 | | (x) |
| HOXD9 | | (x) |

| | | |
|---|---|---|
| HSF1 | (x) | |
| HSF2 | (x) | |
| HTF4 | (x) | |
| HXA1 | (x) | |
| HXA10 | (x) | |
| HXA5 | (x) | |
| HXA7 | (x) | |
| HXB6 | (x) | |
| HXB7 | (x) | |
| HXB8 | (x) | |
| HXC6 | (x) | |
| HXC8 | (x) | |
| HXD10 | (x) | |
| HXD4 | (x) | |
| HXD9 | (x) | |
| IKZF1 | (x) | |
| IRF1 | (x) | |
| IRF2 | (x) | |
| IRF3 | (x) | |
| IRF7 | (x) | (x) |
| IRF9 | (x) | (x) |
| ITF2 | (x) | |
| JUN | (x) | |
| JUNB | (x) | (x) |
| JUND | (x) | (x) |
| KAISO | (x) | |
| KLF1 | (x) | |
| KLF15 | (x) | (x) |
| KLF3 | | (x) |
| KLF4 | (x) | (x) |

| | | |
|---|---|---|
| KLF6 | (x) | (x) |
| KLF8 | (x) | |
| LEF1 | (x) | |
| LYL1 | | (x) |
| MAF | (x) | |
| MAFB | (x) | (x) |
| MAFG | (x) | |
| MAFK.A | (x) | |
| MAFK.S | (x) | |
| MAZ | (x) | (x) |
| MBD2 | (x) | |
| MCR | (x) | |
| MECP2 | (x) | (x) |
| MEF2A | (x) | |
| MEF2C | (x) | |
| MEF2D | (x) | |
| MEIS1 | (x) | |
| MEIS2 | (x) | |
| MITF | (x) | |
| MSX2 | (x) | |
| MTF1 | (x) | |
| MXI1 | (x) | (x) |
| MYB | (x) | (x) |
| MYBB | (x) | |
| MYC | (x) | (x) |
| MYCN | (x) | |
| NANOG.A | (x) | |
| NANOG.S | (x) | |
| NF2L1 | (x) | |
| NF2L2 | (x) | |

| | | |
|---|---|---|
| NFAC1.A | (x) | |
| NFAC1.S | (x) | |
| NFAC3 | (x) | |
| NFAC4 | (x) | |
| NFAT5 | (x) | |
| NFATC3 | | (x) |
| NFATC4 | | (x) |
| NFE2 | (x) | |
| NFE2L1 | | (x) |
| NFIA.C | (x) | |
| NFIA.S | (x) | |
| **NFIB** | | **(x)** |
| NFIL3 | (x) | (x) |
| NFKB1 | (x) | |
| NFKB2 | (x) | (x) |
| NFYA.D | (x) | |
| NFYA.S | (x) | |
| NFYB | (x) | |
| NFYC | (x) | |
| NKX31 | (x) | |
| NR1D1 | (x) | (x) |
| NR1D2 | | (x) |
| NR1H2 | (x) | (x) |
| NR1H3 | | (x) |
| NR1H4 | (x) | |
| NR2C1 | (x) | |
| NR2C2 | (x) | |
| NR2E3 | (x) | |
| NR2F2 | | (x) |
| NR2F6 | (x) | |

| | | |
|---|---|---|
| NR4A1 | | (x) |
| NR4A2 | | (x) |
| NR5A2 | (x) | |
| NR6A1 | (x) | |
| NRF1 | (x) | |
| ONEC2 | (x) | |
| OVOL1 | (x) | (x) |
| P53 | (x) | |
| P63 | (x) | |
| P73 | (x) | |
| PAX5.D | (x) | |
| PBX1 | (x) | |
| PBX2 | (x) | (x) |
| PBX3 | (x) | |
| PEBB | (x) | |
| PIT1 | (x) | |
| PKNX1 | (x) | |
| PLAG1.D | (x) | |
| PLAG1.S | (x) | |
| PO2F1 | (x) | |
| PO2F2 | (x) | |
| PO6F1 | (x) | |
| PPARA.C | (x) | |
| PPARD | (x) | |
| PPARG | | (x) |
| PPARG.A | (x) | |
| PPARG.S | (x) | |
| PRDM1 | (x) | |
| PRGR.C | (x) | |
| PRGR.S | (x) | |

| | | |
|---|---|---|
| PRRX1 | (x) | |
| PRRX2 | (x) | (x) |
| PURA | (x) | |
| RARB | (x) | |
| RARG.C | (x) | |
| RARG.S | (x) | |
| RBPJ | | (x) |
| REL | (x) | |
| RELB | (x) | (x) |
| REST | (x) | |
| RFX1 | (x) | |
| RFX2 | (x) | |
| RFX3 | (x) | |
| RORA | (x) | (x) |
| RORG | (x) | |
| RREB1 | (x) | |
| RUNX1 | (x) | (x) |
| RUNX2 | (x) | (x) |
| RUNX3 | | (x) |
| RXRA | (x) | |
| RXRB | | (x) |
| RXRG | (x) | |
| SHOX2 | | (x) |
| SMAD1 | (x) | |
| SMAD2 | (x) | (x) |
| SMAD3 | | (x) |
| SMAD4 | (x) | |
| SMRC1 | (x) | |
| SNAI1 | (x) | |
| SNAI2 | (x) | (x) |

| | | |
|---|---|---|
| SOX10 | (x) | (x) |
| SOX13 | (x) | |
| SOX17 | (x) | |
| SOX18 | (x) | |
| SOX4 | (x) | (x) |
| SOX5 | (x) | |
| SOX6 | | (x) |
| SOX9 | (x) | |
| SP1.A | (x) | |
| SP1.S | (x) | |
| SP2 | (x) | |
| SP3 | (x) | (x) |
| SP4 | (x) | |
| SPIB | (x) | |
| SRBP1 | (x) | |
| SRBP2 | (x) | |
| SREBF2 | | (x) |
| **STAT5A** | **(x)** | **(x)** |
| **STA5TB** | **(x)** | **(x)** |
| STAT1 | (x) | (x) |
| STAT3 | (x) | |
| STAT4 | (x) | |
| STAT6 | (x) | |
| SUH | (x) | |
| TAL1.A | (x) | |
| TAL1.S | (x) | |
| TBP | (x) | |
| TBX2 | (x) | |
| TBX3 | (x) | (x) |
| TCF3 | | (x) |

| | | |
|---|---|---|
| TCF4 | | (x) |
| TCF7 | (x) | |
| TEAD1 | (x) | |
| TEAD3 | (x) | |
| TEAD4 | (x) | |
| TEF | (x) | |
| TF2L1 | (x) | |
| TF7L2 | (x) | |
| TFAP2A | | (x) |
| TFAP2B | | (x) |
| TFAP2C | | (x) |
| TFCP2 | (x) | |
| TFDP1 | (x) | |
| TFE2.S | (x) | |
| TFE3 | (x) | |
| TFEB | (x) | |
| TGIF1 | | (x) |
| TGIF1.S | (x) | |
| THA.C | (x) | |
| THA.S | (x) | |
| THB.C | (x) | |
| THB.S | (x) | |
| TWST1 | (x) | |
| TYY1 | (x) | |
| USF1 | (x) | |
| USF2 | (x) | |
| VDR | | (x) |
| VDR.C | (x) | |
| VDR.S | (x) | |
| XBP1 | (x) | |

| | | |
|---|---|---|
| YBOX1 | (x) | |
| ZBT18 | (x) | |
| ZBT7A | (x) | |
| ZBTB17 | | (x) |
| ZBTB6 | (x) | |
| ZBTB7A | | (x) |
| ZEB1 | (x) | (x) |
| ZEP1 | (x) | |
| ZEP2 | (x) | |
| ZFHX3 | (x) | |
| ZFP335 | | (x) |
| ZFX | (x) | (x) |
| ZN148 | (x) | |
| ZN423 | (x) | |

**Supplementary Table 5: Comparison of prioritized transcription factors before and after the filtering of the background distribution**

| | Number of TFs before filter | Number of TFs after filter |
|---|---|---|
| TEPIC | 203 | 92 |
| LOG2FC and TEPIC | 203 | 95 |
| TEPIC and DYNAMITE | 203 | 97 |
| DYNAMITE | 203 | 102 |
| LOG2FC and DYNAMITE | 203 | 104 |
| TF-TG score | 203 | 104 |
| LOG2FC | 203 | 119 |

**Supplementary Table 6: Guide which TF was found in which protocol and HM**

| | ATAC-seq | DNase-seq | H2AFZ | H3K27ac | H3K27me3 | H3K36me3 | H3K4me1 | H3K4me2 | H3K4me3 | H3K79me2 | H3K9ac | H3K9me3 | H4K20me1 |
|---|---|---|---|---|---|---|---|---|---|---|---|---|---|
| **AHR** | | | | | | H3K36me3 | | | | | | | |
| **AIRE** | | | H2AFZ | | | | | | | | | | |
| **ALX1** | | | | | | | | H3K4me2 | | | | | |
| **ALX3** | | | | | | | | H3K4me2 | | | | | |
| **ARID5B** | | | H2AFZ | | | | | | | | H3K9ac | | |
| **ARNT** | | | | | | | H3K4me1 | | | | | | |
| **ARNT2** | | | H2AFZ | | | | | | | | | | |
| **ARNT::HIF1A** | ATAC-seq | | | H3K27ac | | | | | | | H3K9ac | | |
| **ASCL1** | | | H2AFZ | | | | | | | | | | |
| **ATF1** | | | | | | | | | | H3K79me2 | | | |
| **ATOH7** | | | | H3K27ac | | | | | | | | | |
| **BACH1** | | DNase-seq | | | | | | | | | | | |

| | | | | | | | | | | | |
|---|---|---|---|---|---|---|---|---|---|---|---|
| BARHL1 | ATAC-seq | | | H3K27ac | | | | | | | |
| BARHL2 | ATAC-seq | | H2AFZ | | | | H3K4me1 | H3K4me2 | | | H3K9ac |
| BARX1 | ATAC-seq | DNase-seq | | | | | H3K4me1 | | | H3K79me2 | |
| BARX2 | ATAC-seq | | | | | | | | | | |
| BATF | ATAC-seq | | | | | | | | | | |
| BATF::JUN | ATAC-seq | | H2AFZ | | | | | | | | H3K9ac |
| BCL11A | | | | | | | | | | | H3K9ac |
| BHLHE22 | | | H2AFZ | | | | | | | | |
| BNC2 | | | | | | | | | | | H3K9ac |
| BSX | ATAC-seq | DNase-seq | H2AFZ | | | | | H3K4me2 | | | H3K9ac |
| CDX2 | | | | H3K27ac | | | | | | | |
| CEBPD | | | | H3K27ac | | | | | | | |
| CLOCK | | | | | | | | H3K4me2 | H3K4me3 | H3K79me2 | |

| | | | | | | | | | | | |
|---|---|---|---|---|---|---|---|---|---|---|---|
| CREB1 | | | | H3K27ac | | | | | | H3K79me2 | H3K9ac |
| CREB3L4 | | | | | | | | H3K4me2 | | | |
| CREM | | | H2AFZ | | | | | | H3K4me3 | | |
| CRX | | | | | | | | H3K4me2 | | | |
| CTCF | | DNase-seq | | | | | | | | | |
| CTCFL | | DNase-seq | | | | | | | | | |
| DLX1 | | | H2AFZ | | | | | | | | H3K9ac |
| DLX3 | | | | | | | | H3K4me2 | | | H3K9ac |
| DLX6 | | | H2AFZ | H3K27ac | | | | | | H3K79me2 | |
| DMRT3 | | DNase-seq | | | | | | | | | |
| DMRTA1 | | | | | | | H3K4me1 | | | | |
| DPRX | ATAC-seq | | | | | H3K36me3 | | | | | |
| DRGX | ATAC-seq | | H2AFZ | | | | | H3K4me2 | | | |

| | | | | | | | | | | | | |
|---|---|---|---|---|---|---|---|---|---|---|---|---|
| E2F5 | | | | | H3K27me3 | | | | H3K4me3 | | | |
| E2F6 | | | | | | | | | | | H3K9ac | |
| E2F8 | | | | H3K27ac | | | | | | | | |
| EBF1 | | | H2AFZ | | | | | | | | | |
| EBF3 | | DNase-seq | | | | | | | | | H3K9ac | H3K9me3 |
| EGR1 | ATAC-seq | | H2AFZ | | | | | | | | H3K9ac | |
| EGR2 | | | H2AFZ | | | | | | | | | |
| EHF | | | H2AFZ | H3K27ac | | | | | | | | |
| ELF1 | | | H2AFZ | | | | H3K4me1 | H3K4me2 | | | | |
| ELF2 | | DNase-seq | | | | | | | H3K4me3 | | | |
| ELF5 | ATAC-seq | DNase-seq | H2AFZ | | | | | | | | | |
| ELK4 | | DNase-seq | | H3K27ac | | | | | | | H3K9ac | |
| EMX1 | | DNase-seq | | | | | H3K4me1 | | | | H3K9ac | |
| EMX2 | ATAC-seq | | | | | | H3K4me1 | | | | | |

| | | | | | | | | | | | |
|---|---|---|---|---|---|---|---|---|---|---|---|
| **EN1** | **ATAC-seq** | **DNase-seq** | **H2AFZ** | | | | | **H3K4me2** | | | **H3K9ac** |
| **EN2** | **ATAC-seq** | | **H2AFZ** | **H3K27ac** | | | **H3K4me1** | | | | **H3K9ac** |
| **EOMES** | | **DNase-seq** | | | | | | | | | |
| **EPAS1** | | | | **H3K27ac** | | | | | | **H3K79me2** | |
| **ERF** | | | | **H3K27ac** | | | | | | | |
| **ERF::FOXI1** | | | | | | | | | | **H3K79me2** | **H3K9ac** |
| **ERF::FOXO1** | | | **H2AFZ** | **H3K27ac** | | | | | | | |
| **ERG** | **ATAC-seq** | | | | | | | **H3K4me2** | | | |
| **ESR2** | | **DNase-seq** | | | | | | | **H3K4me3** | | |
| **ESRRG** | | | | **H3K27ac** | | | | | | | |
| **ESX1** | **ATAC-seq** | **DNase-seq** | | | | | **H3K4me1** | | | | **H3K9ac** |
| **ETS1** | | | | **H3K27ac** | | | | | | | |
| **ETS2** | | | **H2AFZ** | **H3K27ac** | | | | **H3K4me2** | | | |

| | | | | | | | | | | | | |
|---|---|---|---|---|---|---|---|---|---|---|---|---|
| **ETV1** | | | | | | | | **H3K4me2** | | | | |
| **ETV3** | | | | | | | | | | | | **H3K9me3** |
| **ETV4** | | | | **H3K27ac** | | **H3K36me3** | | | | | | |
| **ETV5::DRGX** | | **DNase-seq** | | | | | | | | | | |
| **ETV5::FOXO1** | | | **H2AFZ** | | | | | | | | | |
| **ETV7** | **ATAC-seq** | | **H2AFZ** | | | | | **H3K4me2** | | | | |
| **EVX1** | | **DNase-seq** | | | | | **H3K4me1** | | | | | |
| **EVX2** | | | | | | | | **H3K4me2** | | **H3K79me2** | | |
| **FEZF1** | **ATAC-seq** | | | | | | | | | | | |
| **FIGLA** | | | **H2AFZ** | **H3K27ac** | | | | | | | **H3K9ac** | |
| **FLI1** | | | **H2AFZ** | | | | | | | | | |
| **FOS::JUN** | | | **H2AFZ** | **H3K27ac** | | | | **H3K4me2** | | | | |
| **FOS::JUND** | | **DNase-seq** | | | | | | | | | | |

| | | | | | | | | | | | |
|---|---|---|---|---|---|---|---|---|---|---|---|
| **FOSB** | | | | | | | | | | | **H3K9ac** |
| **FOSB::JUNB** | | | | | | | **H3K4me1** | | | | **H3K9ac** |
| **FOSL1::JUN** | | | | | | | **H3K4me1** | | | | **H3K9ac** |
| **FOSL1::JUNB** | | | | **H3K27ac** | | | | | | | |
| **FOSL1::JUND** | | | **H2AFZ** | **H3K27ac** | | | | | | | |
| **FOSL2::JUN** | | | | | | | | **H3K4me2** | | | **H3K9ac** |
| **FOSL2::JUND** | | | | **H3K27ac** | | | | | | | |
| **FOXA1** | **ATAC-seq** | | | | | | | | | | |
| **FOXA2** | **ATAC-seq** | | **H2AFZ** | | | | | **H3K4me2** | | | **H3K9ac** |
| **FOXA3** | **ATAC-seq** | | **H2AFZ** | | | | | | | | **H3K9ac** |
| **FOXB1** | **ATAC-seq** | **DNase-seq** | | | | | **H3K4me1** | | | | |
| **FOXC1** | **ATAC-seq** | **DNase-seq** | | **H3K27ac** | | | | | | | **H3K9ac** |
| **FOXC2** | **ATAC-seq** | | | | | | | | | **H3K79me2** | |

| | | | | | | | | | | | | |
|---|---|---|---|---|---|---|---|---|---|---|---|---|
| FOXD1 | | | H2AFZ | | | H3K36me3 | | | | | | |
| FOXD3 | ATAC-seq | DNase-seq | H2AFZ | | | | H3K4me1 | | | | | |
| FOXF2 | | DNase-seq | | | | | | | | H3K79me2 | | |
| FOXG1 | | DNase-seq | | | | | | | | | | |
| FOXI1 | | DNase-seq | | | | H3K36me3 | | | | | H3K9ac | |
| FOXJ2 | ATAC-seq | | | | | | | | | | H3K9ac | |
| FOXJ2::ELF1 | | DNase-seq | H2AFZ | | | | | | H3K4me3 | | | |
| FOXK1 | | DNase-seq | | | | | H3K4me1 | | | | | |
| FOXK2 | | | H2AFZ | | | | | | | H3K79me2 | | |
| FOXL1 | | | | | | | | H3K4me2 | | | | |
| FOXM1 | | | | | | | | | | H3K79me2 | | |
| FOXN3 | | DNase-seq | | | | | | | | | | |
| FOXO3 | | DNase-seq | | | | | | | | | | H3K9me3 |

| | | | | | | | | | | | |
|---|---|---|---|---|---|---|---|---|---|---|---|
| **FOXO4** | | **DNase-seq** | | | | | **H3K4me1** | **H3K4me2** | | | |
| **FOXO6** | **ATAC-seq** | **DNase-seq** | | | | | | | | | **H3K9ac** |
| **FOXP1** | **ATAC-seq** | | **H2AFZ** | | | | | | | **H3K79me2** | |
| **FOXP2** | **ATAC-seq** | **DNase-seq** | | | | | | | | | |
| **FOXP3** | | | | | | | | | | **H3K79me2** | |
| **FOXQ1** | | | | | | | **H3K4me1** | **H3K4me2** | | | |
| **GABPA** | | | **H2AFZ** | | | | **H3K4me1** | | | | **H3K9ac** |
| **GATA1** | | **DNase-seq** | | | | | | | | | |
| **GATA2** | | | | | | | | | | **H3K79me2** | |
| **GATA3** | **ATAC-seq** | | | **H3K27ac** | | | **H3K4me1** | | | | **H3K9ac** |
| **GATA4** | **ATAC-seq** | | | **H3K27ac** | | | | | **H3K4me3** | | **H3K9ac** |
| **GATA5** | **ATAC-seq** | | | | **H3K27me3** | | | | | | |
| **GATA6** | | | | | | | | **H3K4me2** | | | **H3K9ac** |

| | | | | | | | | | | | |
|---|---|---|---|---|---|---|---|---|---|---|---|
| **GBX1** | **ATAC-seq** | **DNase-seq** | | | | | **H3K4me1** | | | | |
| **GBX2** | | **DNase-seq** | | | | **H3K36me3** | | **H3K4me2** | | | **H3K9ac** |
| **GCM2** | | | | | | | | | | **H3K79me2** | **H3K9ac** |
| **GMEB2** | | | **H2AFZ** | | | | | | | | |
| **GRHL1** | | | | | **H3K27me3** | | | | | | |
| **GRHL2** | | | **H2AFZ** | | | | | | | | |
| **GSC** | **ATAC-seq** | | | **H3K27ac** | | | | | | | |
| **GSC2** | | | | **H3K27ac** | | | **H3K4me1** | | | | |
| **GSX2** | | **DNase-seq** | | | | | | **H3K4me2** | | | **H3K9ac** |
| **HES2** | | | **H2AFZ** | | | | | | | | |
| **HES5** | | **DNase-seq** | **H2AFZ** | | | | | | | | |
| **HES6** | | **DNase-seq** | | | | | | | | | |
| **HESX1** | **ATAC-seq** | **DNase-seq** | **H2AFZ** | **H3K27ac** | | | | | | | **H3K9ac** |
| **HEY1** | | **DNase-seq** | | | | | | | | | **H3K9ac** |

| | | | | | | | | | | | |
|---|---|---|---|---|---|---|---|---|---|---|---|
| HEY2 | ATAC-seq | DNase-seq | | | | | | | | | |
| HIC1 | | | | | | | H3K4me1 | H3K4me2 | | | H3K9ac |
| HIC2 | | | | | | | | | H3K4me3 | | |
| HIF1A | | | | | | | | | H3K4me3 | | |
| HINFP | | DNase-seq | | | | | | | | | |
| HMBOX1 | | | | | | H3K36me3 | H3K4me1 | | | | |
| HNF4G | | | H2AFZ | | | | H3K4me1 | | | | |
| HOXA1 | | | | | | | | | | H3K79me2 | H3K9ac |
| HOXA10 | ATAC-seq | DNase-seq | | | | | | H3K4me2 | | | |
| HOXA2 | | DNase-seq | | H3K27ac | | | | | | | |
| HOXA4 | | | | | | | | | | | H3K9ac |
| HOXA6 | | | | | | | H3K4me1 | H3K4me2 | | | H3K9ac |
| HOXA7 | | | | | | | | H3K4me2 | | | |

| | | | | | | | | | | | | | |
|---|---|---|---|---|---|---|---|---|---|---|---|---|---|
| HOXA9 | | | H2AFZ | | | | | | H3K4me3 | | | | |
| HOXB2 | ATAC-seq | DNase-seq | H2AFZ | | | H3K36me3 | | H3K4me2 | | H3K79me2 | | | |
| HOXB3 | | | | | | | | H3K4me2 | | H3K79me2 | | | |
| HOXB4 | | | | | | | | H3K4me2 | | | | | |
| HOXB5 | | | | | | | H3K4me1 | H3K4me2 | | | | | |
| HOXB6 | | | H2AFZ | H3K27ac | | | | | | | | | |
| HOXB7 | | | H2AFZ | | | | | | | | | | |
| HOXB8 | ATAC-seq | | H2AFZ | | | | | | | | | | H4K20me1 |
| HOXC10 | ATAC-seq | | | | | | | | | | | | |
| HOXC4 | ATAC-seq | DNase-seq | H2AFZ | | | | | | | H3K79me2 | | | |
| HOXC8 | | | | | | | | H3K4me2 | H3K4me3 | | | | |
| HOXD11 | | DNase-seq | | | | | | | | | | | |
| HOXD4 | | | H2AFZ | | | | | | | | H3K9ac | | |
| HOXD8 | ATAC-s | | H2AFZ | | | | | H3K4m | | H3K79 | | | H4K20 |

| | | | | | | | | | | | | | |
|---|---|---|---|---|---|---|---|---|---|---|---|---|---|
| | eq | | | | | | | e2 | | me2 | | | me1 |
| HOXD9 | ATAC-seq | | | | | | | | | | | | |
| IKZF1 | | | | | | | | H3K4me2 | | | | | |
| IRF2 | | | H2AFZ | | | | | | | | | | |
| ISX | | | | H3K27ac | | | | | | H3K79me2 | | | |
| JUN::JUNB | | | | | | | | H3K4me2 | | H3K79me2 | | | |
| JUND | ATAC-seq | | | | | | | | | | | | |
| KLF1 | | | | H3K27ac | | | | | | | H3K9ac | | |
| KLF10 | ATAC-seq | DNase-seq | | H3K27ac | | | | | | | | | |
| KLF11 | | | | | | | | | | H3K79me2 | | | |
| KLF12 | | DNase-seq | | | | | H3K4me1 | H3K4me2 | | H3K79me2 | | | |
| KLF14 | | | | | | | H3K4me1 | H3K4me2 | | | | | |
| KLF15 | | | H2AFZ | | | | | | | | | | |
| KLF16 | | | H2AFZ | | | | | | | | H3K9ac | | |

| | | | | | | | | | | | |
|---|---|---|---|---|---|---|---|---|---|---|---|
| **KLF2** | | | **H2AFZ** | | | **H3K36me3** | | | | | **H3K9ac** |
| **KLF3** | | | | | | | | **H3K4me2** | | | |
| **KLF4** | | | **H2AFZ** | | | | **H3K4me1** | | | **H3K79me2** | |
| **KLF6** | | | | | | | **H3K4me1** | **H3K4me2** | | | |
| **KLF7** | | | | | | | **H3K4me1** | **H3K4me2** | | | |
| **KLF8** | | | **H2AFZ** | | | | | | | | |
| **LBX2** | | | | | | | **H3K4me1** | **H3K4me2** | | **H3K79me2** | |
| **LHX1** | | | | | | **H3K36me3** | | | | | **H3K9ac** |
| **LHX2** | | | | | | | **H3K4me1** | | | | |
| **LHX5** | | | | **H3K27ac** | | | | | | | |
| **MAFB** | | | | | | | | | **H3K4me3** | | |
| **MAFG** | | **DNase-seq** | | | **H3K27me3** | | | | | | |
| **MAFK** | | | **H2AFZ** | | | | | | | | |
| **MAX** | | | **H2AFZ** | | | | | | | | **H3K9ac** |

| | | | | | | | | | | | |
|---|---|---|---|---|---|---|---|---|---|---|---|
| **MAX::MYC** | **ATAC-seq** | | | | | | | **H3K4me2** | | | |
| **MAZ** | | | **H2AFZ** | **H3K27ac** | | | | | | | **H3K9ac** |
| **MECP2** | | | | | | | **H3K4me1** | | | | |
| **MEF2A** | | **DNase-seq** | | **H3K27ac** | | | | **H3K4me2** | | | |
| **MEF2C** | | | | | | | | | | **H3K79me2** | |
| **MEIS1** | **ATAC-seq** | | | | | | | | **H3K4me3** | | |
| **MEIS2** | **ATAC-seq** | | | | | | | | | | |
| **MEIS3** | **ATAC-seq** | | | | | | | | | | |
| **MEOX1** | | **DNase-seq** | **H2AFZ** | | | | **H3K4me1** | | | | **H3K9ac** |
| **MEOX2** | | | | | | | | | | **H3K79me2** | |
| **MGA** | | **DNase-seq** | **H2AFZ** | **H3K27ac** | | | | | | | |
| **MGA::EVX1** | | **DNase-seq** | | | | | | | | | |
| **MIXL1** | | | **H2AFZ** | | | | | | | **H3K79me2** | |

| | | | | | | | | | | | | |
|---|---|---|---|---|---|---|---|---|---|---|---|---|
| **MNT** | | **DNase-seq** | **H2AFZ** | **H3K27ac** | | | | **H3K4me2** | **H3K4me3** | **H3K79me2** | | |
| **MNX1** | **ATAC-seq** | | **H2AFZ** | | | | | **H3K4me2** | | | **H3K9ac** | |
| **MSANTD3** | **ATAC-seq** | **DNase-seq** | | **H3K27ac** | | | | | | | | |
| **MSGN1** | **ATAC-seq** | | | | | | | | | **H3K79me2** | **H3K9ac** | |
| **MSX1** | | | | | | | **H3K4me1** | | | | **H3K9ac** | |
| **MSX2** | **ATAC-seq** | | | **H3K27ac** | | | | | | **H3K79me2** | | |
| **MXI1** | | | **H2AFZ** | | | | | **H3K4me2** | | | **H3K9ac** | |
| **MYB** | | | **H2AFZ** | **H3K27ac** | | | | **H3K4me2** | | | | |
| **MYC** | | | | | | | | | | **H3K79me2** | | |
| **MYF5** | | | | | | | | | **H3K4me3** | | | |
| **MYOD1** | | | **H2AFZ** | **H3K27ac** | | | | **H3K4me2** | **H3K4me3** | | | |
| **MYOG** | | | | **H3K27ac** | | | | | | | | |
| **MZF1** | | | | | | | | | | | | **H3K9me3** |

| | | | | | | | | | | | |
|---|---|---|---|---|---|---|---|---|---|---|---|
| NEUROD1 | | | H2AFZ | | | | | | | | H3K9ac |
| NEUROG2 | | | H2AFZ | H3K27ac | | | | H3K4me2 | | | |
| NFATC1 | | | | | | | H3K4me1 | | | | |
| NFATC4 | ATAC-seq | DNase-seq | | H3K27ac | | | | | | | H3K9ac |
| NFE2L1 | | | | H3K27ac | | | | H3K4me2 | | | H3K9ac |
| NFE2L2 | | | | H3K27ac | | | | | | | |
| NFIL3 | | | | | | | | | | H3K79me2 | |
| NFYA | | | | H3K27ac | | | | H3K4me2 | H3K4me3 | | |
| NFYB | | | H2AFZ | | | | | | | | |
| NFYC | | | | | | | | | | H3K79me2 | |
| NHLH1 | | | H2AFZ | | | H3K36me3 | | | | | |
| NOBOX | | | H2AFZ | | | | | H3K4me2 | | | |
| NOTO | ATAC-seq | DNase-seq | H2AFZ | H3K27ac | | | H3K4me1 | | | H3K79me2 | H3K9ac |

| | | | | | | | | | | | | |
|---|---|---|---|---|---|---|---|---|---|---|---|---|
| **NR1H3** | | | | | | | | | **H3K4me3** | | | |
| **NR1H4** | | | | **H3K27ac** | | | | | | | | |
| **NR1I3** | | | **H2AFZ** | | | | | | | | | |
| **NR2C1** | **ATAC-seq** | | | **H3K27ac** | | | | | | | | |
| **NR2C2** | | | **H2AFZ** | | | | | | | | **H3K9ac** | |
| **NR2F2** | | **DNase-seq** | | | | | | | | **H3K79me2** | | |
| **NR4A1** | | **DNase-seq** | **H2AFZ** | | | **H3K36me3** | **H3K4me1** | | | | | **H3K9me3** |
| **NR4A2** | | **DNase-seq** | | | | | | | | | | |
| **NR5A1** | | | | | | | **H3K4me1** | | | | | |
| **NR5A2** | | | **H2AFZ** | | | | | **H3K4me2** | | | | |
| **NRF1** | | | **H2AFZ** | | | | | | | | | |
| **OSR1** | | | **H2AFZ** | | | | | | **H3K4me3** | | | |
| **OSR2** | | | | | | | | **H3K4me2** | | | | |
| **OTX1** | | | **H2AFZ** | | | | | | | | | |

| | | | | | | | | | | | |
|---|---|---|---|---|---|---|---|---|---|---|---|
| **OTX2** | | | | | | | | **H3K4me2** | | **H3K79me2** | **H3K9ac** |
| **OVOL2** | | | | | **H3K27me3** | | | | | | |
| **PATZ1** | | | **H2AFZ** | **H3K27ac** | | | | | | | **H3K9ac** |
| **PAX4** | | **DNase-seq** | | | | | | | | | **H3K9ac** |
| **PAX5** | **ATAC-seq** | | **H2AFZ** | | | | | | | | |
| **PBX1** | | | **H2AFZ** | | | | | | | | |
| **PDX1** | **ATAC-seq** | **DNase-seq** | | | | | | **H3K4me2** | | | **H3K9ac** |
| **PGR** | | | | | | | | | **H3K4me3** | | |
| **PITX1** | **ATAC-seq** | | | | | | | **H3K4me2** | | | |
| **PITX3** | | | | | | | | | | | **H3K9ac** |
| **PLAG1** | | | | | | **H3K36me3** | | **H3K4me2** | **H3K4me3** | | |
| **PLAGL2** | | | | | | | | **H3K4me2** | | | |
| **POU2F1** | | **DNase-seq** | | | | | | | | | **H3K9ac** |
| **POU3F** | | | | | | | **H3K4m** | | | **H3K79** | |

| | | | | | | | | | | | |
|---|---|---|---|---|---|---|---|---|---|---|---|
| 1 | | | | | | | e1 | | | me2 | |
| POU3F3 | | DNase-seq | | | | | | | | | |
| POU5F1 | | | | H3K27ac | | H3K36me3 | | | | | |
| POU5F1B | | | H2AFZ | | | | | | | | |
| POU6F1 | | DNase-seq | H2AFZ | | | | | | | | |
| POU6F2 | | | | | | | | | | H3K79me2 | |
| PPARA | | | H2AFZ | | | | | | H3K4me3 | | |
| PRDM14 | | | | | | | | | H3K4me3 | | |
| PRDM4 | | | H2AFZ | | | | | H3K4me2 | | | |
| PRDM6 | ATAC-seq | | H2AFZ | H3K27ac | | | | | | | H3K9ac |
| PRRX1 | | DNase-seq | H2AFZ | | | | H3K4me1 | | | | |
| PRRX2 | | DNase-seq | | H3K27ac | | | H3K4me1 | | | | |
| RARA::RXRG | | DNase-seq | H2AFZ | | | | | | H3K4me3 | | |

| | | | | | | | | | | | |
|---|---|---|---|---|---|---|---|---|---|---|---|
| RAX | ATAC-seq | | H2AFZ | | | | | | H3K4me3 | H3K79me2 | |
| RAX2 | | | | | | | | H3K4me2 | | H3K79me2 | H3K9ac |
| RBPJ | | | | H3K27ac | | | | H3K4me2 | | | H3K9ac |
| REL | | | | | | | | | | H3K79me2 | H3K9ac |
| RELA | ATAC-seq | | H2AFZ | | | | | H3K4me2 | | | |
| RELB | | | H2AFZ | | | | | H3K4me2 | | | |
| RFX7 | | | | | | | | | | | H3K9ac |
| RHOXF1 | | | | | | | | | H3K4me3 | | |
| RORA | | | H2AFZ | H3K27ac | | | | H3K4me2 | | | |
| RORC | | | | | | | | H3K4me2 | | H3K79me2 | H3K9ac |
| RUNX1 | | | | | | | | H3K4me2 | H3K4me3 | | |
| RUNX3 | | | H2AFZ | | | | | H3K4me2 | | | |
| RXRA | ATAC-seq | | | | | | | | | | |

| | | | | | | | | | | | |
|---|---|---|---|---|---|---|---|---|---|---|---|
| **SALL4** | | | **H2AFZ** | | | | **H3K4me1** | **H3K4me2** | | **H3K79me2** | **H3K9ac** |
| **SATB1** | | **DNase-seq** | | | | | | | | | |
| **SCRT1** | | **DNase-seq** | | | | | | | | | |
| **SCRT2** | | **DNase-seq** | | | | | | | | | |
| **SIX1** | | **DNase-seq** | | | | **H3K36me3** | | | | | |
| **SIX2** | | **DNase-seq** | | | | | | | | | |
| **SMAD2** | | | | | | | | | **H3K4me3** | | |
| **SMAD4** | | | **H2AFZ** | | | | | | | | |
| **SNAI1** | **ATAC-seq** | | | | | **H3K36me3** | | | | | |
| **SNAI2** | | **DNase-seq** | | | | | | | | | **H3K9ac** |
| **SNAI3** | | | | | | **H3K36me3** | | | | | |
| **SOHLH2** | | | | **H3K27ac** | | | | | | | |
| **SOX10** | **ATAC-seq** | | **H2AFZ** | | | | | | | **H3K79me2** | |

| | | | | | | | | | | | |
|---|---|---|---|---|---|---|---|---|---|---|---|
| SOX13 | | | | | | | H3K4me1 | H3K4me2 | | | |
| SOX15 | ATAC-seq | DNase-seq | | | | | | H3K4me2 | | H3K79me2 | |
| SOX17 | ATAC-seq | | | | | | H3K4me1 | | | | |
| SOX2 | | | | H3K27ac | | | | | | H3K79me2 | H3K9ac |
| SOX3 | ATAC-seq | DNase-seq | | | | H3K36me3 | | H3K4me2 | | | |
| SOX4 | ATAC-seq | | H2AFZ | | | | | | | | H3K9ac |
| SOX8 | ATAC-seq | DNase-seq | H2AFZ | | | | H3K4me1 | | | H3K79me2 | |
| SP1 | | | H2AFZ | | | | | | | | H3K9ac |
| SP2 | ATAC-seq | | | | | | | | | | |
| SP3 | | DNase-seq | | | | | | | | | |
| SP4 | | | | | | | | | | | H3K9ac |
| SP8 | | | H2AFZ | H3K27ac | | | | | | | |
| SP9 | | | H2AFZ | | | | | H3K4me2 | | H3K79me2 | |
| SREBF | | | H2AFZ | H3K27a | | | | | | | |

| | | | | | | | | | | | | |
|---|---|---|---|---|---|---|---|---|---|---|---|---|
| 1 | | | | c | | | | | | | | |
| SREBF2 | | | | | | | | H3K4me2 | | | | H3K9me3 |
| SRY | | DNase-seq | | | | | | | | | | |
| STAT3 | ATAC-seq | | | | | | H3K4me1 | | | | | |
| STAT5A | | | | H3K27ac | | | | | | | | |
| STAT5B | ATAC-seq | | | | | | | | | | | |
| STAT6 | | | H2AFZ | | | H3K36me3 | | | | | | |
| TAF1 | | DNase-seq | | H3K27ac | | | | | | H3K79me2 | | |
| TAL1::TCF3 | | DNase-seq | | | | | | H3K4me2 | | | | |
| TBP | | | | | | H3K36me3 | | | | | | |
| TBX1 | | | | H3K27ac | | | | | | | | |
| TBX18 | | | H2AFZ | | | | | | | | H3K9ac | |
| TBX5 | | | H2AFZ | H3K27ac | | | | | | | | |
| TBX6 | | | | | | | | H3K4m | | | | |

| | | | | | | | | | | | |
|---|---|---|---|---|---|---|---|---|---|---|---|
| | | | | | | | | e2 | | | |
| TCF12 | | | H2AFZ | | | | | | | | H3K9ac |
| TCF21 | | | | | | H3K36me3 | | | | | |
| TCF3 | | | H2AFZ | | | | H3K4me1 | | | | |
| TCF4 | | | | | | | | H3K4me2 | | H3K79me2 | |
| TCF7 | | | H2AFZ | | | | | | | | |
| TCF7L2 | ATAC-seq | DNase-seq | | | | | | | | | |
| TCFL5 | | | | | | | | | H3K4me3 | | H3K9ac |
| TEAD2 | | | | | | | | | | | H3K9ac |
| TEAD4 | | | H2AFZ | H3K27ac | | | H3K4me1 | H3K4me2 | | | |
| TFAP2A | | | | | | | | | | H3K79me2 | |
| TFAP2B | | | H2AFZ | H3K27ac | | | | H3K4me2 | | | |
| TFAP2C | | | H2AFZ | | | | | | | | H3K9ac |
| TFAP4 | | DNase-seq | | | | | | | | | |

| | | | | | | | | | | | |
|---|---|---|---|---|---|---|---|---|---|---|---|
| **TFDP1** | | | **H2AFZ** | | | | | | | | **H3K9ac** |
| **TFEB** | | | **H2AFZ** | | | | | | | | **H3K9ac** |
| **THAP1** | | | | | | **H3K36 me3** | | | | | |
| **TLX2** | **ATAC-s eq** | **DNase-seq** | | | | | | **H3K4m e2** | | | |
| **TRPS1** | | | **H2AFZ** | | | | **H3K4m e1** | | | | |
| **TWIST1** | | **DNase-seq** | | | | | | | | | **H3K9ac** |
| **VAX1** | | | | | | | | | | **H3K79 me2** | **H3K9ac** |
| **VAX2** | **ATAC-s eq** | **DNase-seq** | **H2AFZ** | | | | | **H3K4m e2** | | | **H3K9ac** |
| **VENTX** | **ATAC-s eq** | **DNase-seq** | | **H3K27a c** | | | | | **H3K4m e3** | **H3K79 me2** | **H3K9ac** |
| **VSX1** | | | **H2AFZ** | | | | | | | | |
| **VSX2** | | | **H2AFZ** | | | | | **H3K4m e2** | | **H3K79 me2** | **H3K9ac** |
| **WT1** | | **DNase-seq** | **H2AFZ** | | | | | | | | **H3K9ac** |
| **YY2** | | | **H2AFZ** | | | | | | | | |
| **ZBED2** | **ATAC-s eq** | | | | | | | | | | **H3K9ac** |

| | | | | | | | | | | | |
|---|---|---|---|---|---|---|---|---|---|---|---|
| **ZBTB17** | | **DNase-seq** | | | | | | | **H3K4me3** | | |
| **ZBTB26** | | | | **H3K27ac** | | | | **H3K4me2** | | | |
| **ZBTB6** | | | **H2AFZ** | | | | | **H3K4me2** | | | |
| **ZBTB7A** | | | | **H3K27ac** | | | | | | | |
| **ZBTB7B** | | **DNase-seq** | | | | | | | | | |
| **ZBTB7C** | **ATAC-seq** | **DNase-seq** | | | | | | | | **H3K79me2** | |
| **ZFP14** | | | **H2AFZ** | | | | | | | | |
| **ZFP57** | | | | | | | | | | | **H3K9ac** |
| **ZFX** | | | | | | | | | **H3K4me3** | | |
| **ZIC4** | | **DNase-seq** | **H2AFZ** | **H3K27ac** | | | | **H3K4me2** | | | |
| **ZKSCAN5** | | | | | | **H3K36me3** | | | | | |
| **ZNF148** | | | | | | | | | | **H3K79me2** | |
| **ZNF18** | | | **H2AFZ** | | | | | | | | |
| **ZNF189** | | | **H2AFZ** | | | | | **H3K4me2** | | | |

| | | | | | | | | | | | |
|---|---|---|---|---|---|---|---|---|---|---|---|
| ZNF263 | | | | | | | | H3K4me2 | | | |
| ZNF317 | | | H2AFZ | | | | | | | | |
| ZNF331 | | | H2AFZ | | | | | | | | |
| ZNF350 | ATAC-seq | | H2AFZ | H3K27ac | | | | | H3K4me3 | | |
| ZNF354A | | | | | | | H3K4me1 | | | | |
| ZNF354C | | | H2AFZ | | | | | | | | |
| ZNF384 | ATAC-seq | | | | | | H3K4me1 | H3K4me2 | | | |
| ZNF394 | | | | | | | | | | | H3K9ac |
| ZNF454 | | | H2AFZ | H3K27ac | | | | | | | H3K9ac |
| ZNF460 | | | | | | | | | | H3K79me2 | |
| ZNF530 | | | H2AFZ | | | | | | | | |
| ZNF549 | | | H2AFZ | | | | | H3K4me2 | | | |
| ZNF554 | | DNase-seq | | | | | | | | | H3K9ac |
| ZNF563 | ATAC-seq | | H2AFZ | | | H3K36me3 | | | H3K4me3 | | |

| | | | | | | | | | | | | |
|---|---|---|---|---|---|---|---|---|---|---|---|---|
| **ZNF610** | | | **H2AFZ** | **H3K27ac** | | | | | | | | |
| **ZNF652** | | | | | | | | | | | | **H3K9me3** |
| **ZNF667** | | | | | | | | | **H3K4me3** | | | |
| **ZNF682** | | **DNase-seq** | | | | | | | | | | |
| **ZNF692** | | | | **H3K27ac** | | | | | | | | |
| **ZNF740** | | | **H2AFZ** | | | | | | | | | |
| **ZNF770** | | | **H2AFZ** | | | | | | | | | |
| **ZNF784** | | **DNase-seq** | | | | | | | | | | |
| **ZNF93** | | **DNase-seq** | | | | | | | | | | |

14. Thorvaldsdóttir H, Robinson JT, Mesirov JP. Integrative Genomics Viewer (IGV): high-performance genomics data visualization and exploration. Brief Bioinform.

2013;14: 178–192.

22. Ghosh, Ezashi, Ostrowski. A central role for Ets-2 in the transcriptional regulation and cyclic adenosine 5′-monophosphate responsiveness of the human chorionic gonadotropin-β subunit …. Mol Life. Available: https://academic.oup.com/mend/article-abstract/17/1/11/2741656

38. National Center for Biotechnology Information (US). The p53 tumor suppressor protein. National Center for Biotechnology Information (US); 1998.
